# Different Coexisting Mpox Lineages Were Continuously Circulating in Humans Prior to 2022

**DOI:** 10.1101/2023.01.03.522633

**Authors:** Nnaemeka Ndodo, Jonathan Ashcroft, Kuiama Lewandowski, Adesola Yinka-Ogunleye, Chimaobi Chukwu, Adama Ahmad, David King, Afolabi Akinpelu, Carlos Maluquer de Motes, Paolo Ribeca, Rebecca P. Sumner, Andrew Rambaut, Michael Chester, Tom Maishman, Oluwafemi Bamidele, Nwando Mba, Olajumoke Babatunde, Olusola Aruna, Steven T. Pullan, Benedict Gannon, Colin Brown, Chikwe Ihekweazu, Ifedayo Adetifa, David O. Ulaeto

## Abstract

The origin and hazardous potential of human mpox is obscured by a lack of genomic data between the 2018, when exportations from Nigeria were recorded, and 2022 when the global outbreak started. Here, 18 genomes from patients across southern Nigeria in 2019/20 reveal multiple lineages of Monkeypox virus have achieved sustained human-to-human transmission, co-existing in humans for several years and accumulating mutations consistent with APOBEC3 activity suggesting the virus in humans is now segregated from its natural reservoir. Remarkably, three genomes have disruptions in the A46R gene, which contributes to innate immune modulation. The data demonstrates that the A.2 lineage, multiply exported to North America since 2021 independently of the global outbreak, has persisted in Nigeria for more than two years prior to its latest exportation.

**One-Sentence Summary:** Mpox is now a human diseae evolving in humans with multiple variants taking separate paths towards adaptation, some analogous to those of *Variola*

## Main Text

*Monkeypox virus* (MPXV) is an orthopoxvirus (OPV) endemic to West and Central Africa, circulating in one or more rodent species with epizootic infections of monkeys and chimpanzees, and zoonotic infections of humans. Human monkeypox, now known as Mpox, is traditionally a rash illness similar to smallpox with inefficient human-to-human transmission (*1*). An ongoing human outbreak was reported from 2017 in Nigeria, with tens of confirmed cases per year (*2-4*). As a result, a few cases of Mpox were detected between 2018 and 2021 in individuals travelling from Nigeria to non-endemic countries (*4-8*). In 2022 an ongoing global outbreak occurred in individuals with no history of travel to Africa, with extended human-to-human transmission, concentrated among men-who-have-sex-with-men (MSM) communities (*7, 9*), indicating the virus has found a novel route to transmit between humans without re-introduction from animal reservoirs. As a zoonotic virus successfully adopting non-zoonotic transmission between humans, MPXV is unlikely to be optimally adapted to survival and transmission across its new host. Instead adaptive mutations are to be expected along with dispersal and divergence driven by founder effects and selection pressure (*10*). Initial analysis of several publicly available genome sequences for human MPXV isolates from 2017 onwards indicate that the 2022 global MPXV outbreak (now known as lineage B.1) is phylogenetically linked with the 2017 Nigerian outbreak and most likely has a single origin (*11*). The evolutionary path resulting in the genesis of lineage B.1 remains obscure with few travel-associated isolates available between 2018 and 2022. In addition, a second lineage has recently been described from 3 cases introduced in the United States between 2021 and 2022 (*12*). Here, we have analysed 18 MPXV genomes from patients in Nigeria in 2019 and 2020, and possibly the only Clade II human mpox genomes available from this period. A phylogenetic tree demonstrates the 18 sequences are from clade IIb (*13, 14*) and cluster within lineage A (Figure 1A). A total of 187 polymorphisms, 149 of which are single nucleotide polymorphisms (SNPs) and 35 of which are common to 17 of the genomes, were identified relative to the 1971 KJ642617 reference sequence (*15, 16*) (Table S1). This suggests mutation and potentially adaption in infected people, although heterogeneity in circulating MPXV in the reservoir host is not excluded, and assuming the outbreak stems from a single zoonotic crossover. The identified polymorphisms are distributed throughout the genome and many (84) are associated with alterations to protein primary sequence (Table S1). They include both uncharacterised genes and well characterised genes/proteins, including the virus F13 membrane protein (two instances of synonymous SNP) and the E9 DNA polymerase (two instances of non-synonymous missense SNP), which are the targets of the licensed smallpox antivirals Tecovirimat and Brincidofovir respectively. It is not expected that efficacy of the drugs will be affected, given that neither was in use in Nigeria at the time, and their broad activity for OPVs. However, it raises the possibility that sub-optimal deployment of these drugs could drive evolution of the target genes. The 18 genomes described here came from patient samples taken between January 2019 and January 2020, widely distributed across the southern part of Nigeria (Figure 1B). Available sequences from the lineage B.1 form a distinct branch from this cluster (Figure 1A). This suggests that a founder effect may have operated in the genesis of the international outbreak and that lineage B.1 should therefore have a lower baseline variation than the wider Clade II population that has been causing human infections in Nigeria since at least 2017. Altogether our analysis demonstrates the existence of divergent variants circulating in Nigeria before the emergence of lineage B.1.

**Fig. 1.**
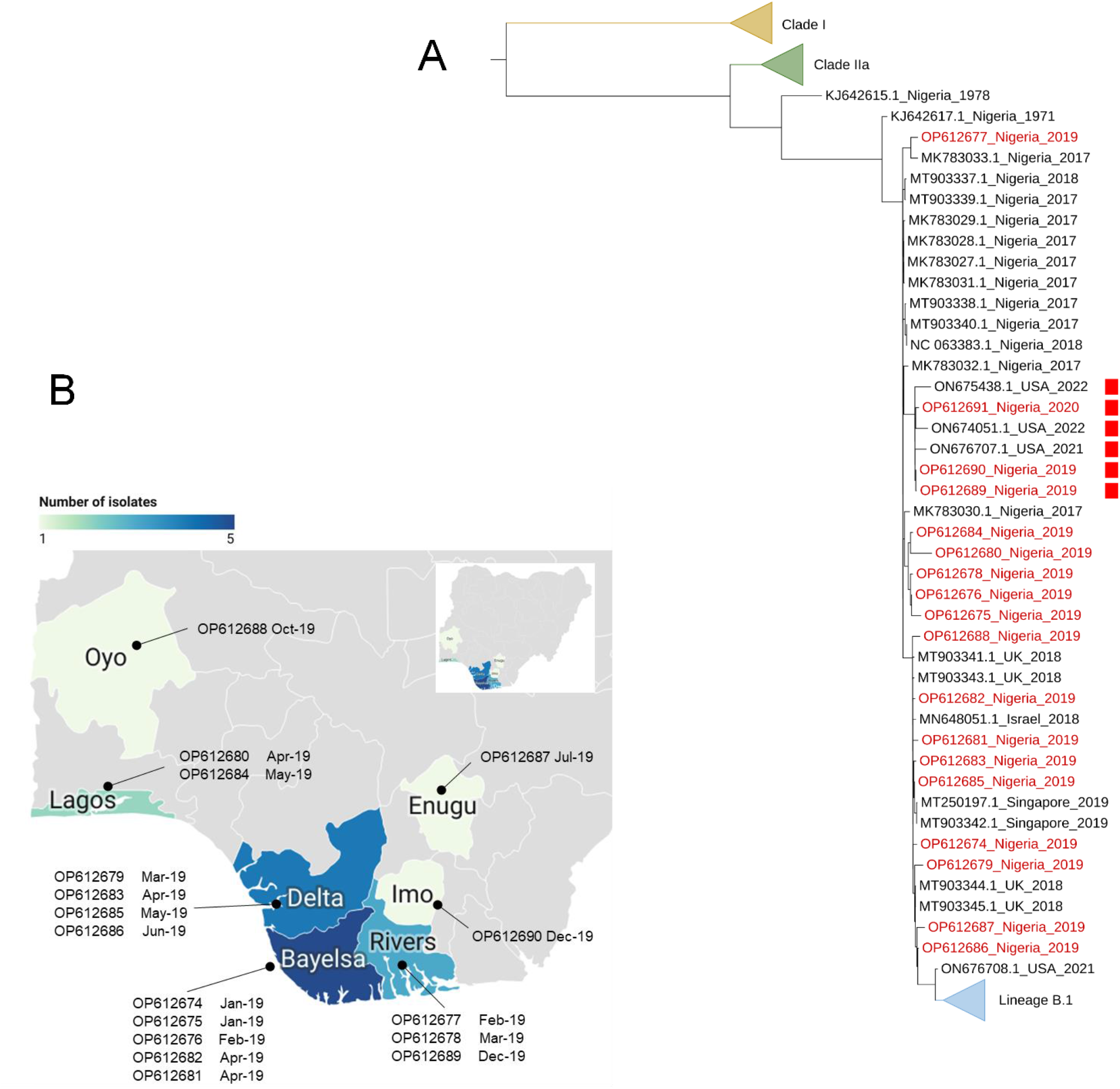
A) Phylogenetic Tree of Clade II MPXV genomes. Genomes described in this study are highlighted in red and those with a disrupted A46R gene are marked with a red square. Published genomes from the current international outbreak are depicted in a single cluster in grey. The cut off date for inclusion of newly published genomes was June 18th 2022. B) Expanded map showing distribution of cases from which the genomes described in the study were isolated, across southern states of Nigeria.

### Mutations in putative APOBEC3 motifs

Analysis of multiple genomes from the 2022 international outbreak has identified 47 shared single nucleotide mutations relative to the 2018 UK reference genome MT903345 (*11, 17*). Of these, 45 were compatible with the action of APOBEC3 cytidine deaminase activity (*17*). Using the same reference genome for a baseline, all 18 of the sequences described here have APOBEC3 style mutations, but at a lower frequency than in the 2022 sequences from lineage B.1 (Figure 2A). OP612686 and OP612687 share the most APOBEC3 style mutations with genomes from the 2022 B.1 lineage and are also closest to these sequences on the phylogenetic tree. This indicates that sustained non-zoonotic transmission is facilitating MPXV acquiring APOBEC3 style mutations, and it has been suggested this could provide the basis for a molecular clock to calculate the origin of the initial zoonotic crossover (*18*). There is considerable variation among the 18 genomes described here with respect to APOBEC3 style mutations, with the mutations occurring at different places in different genomes (Figure 3 and Table S1). The frequency of APOBEC3 style mutations relative to MT903345 (median 8.5) is much lower than the frequency in genomes from Lineage B.1 (median 42). This difference is statistically significant (*P* = 8.37×10^−14^, Kruskall-Wallis test) (Table S2). The elevated APOBEC3 style mutations observed in Lineage B.1 sequences relative to those described here could be a function of random accumulation over time, or alternatively reflect APOBEC3 enabled adaptation during the international outbreak. Repeating the analysis as a comparison of the 18 genomes, and MT903345, against the 1971 KJ642617 genome demonstrates a higher frequency of APOBEC3 style mutations in the 18 genomes (median 25), and a significant frequency in MT903345 and other genomes isolated in 2018 (*4, 19*) (median 22.5). The median frequency of APOBEC3 style mutations in Lineage B.1 genomes rises to 67 when compared with KJ642617 (Figure 2B) (Table S3). Although there is a year-on-year increase in APOBEC3 style mutations from 2017 onwards, the only statistically significant differences are between 2022 and the other years. Pairwise comparisons between years 2017 to 2021 are not significant (Table S4). This raises the possibility the higher frequency of APOBEC3 style mutations in the B.1 2022 temporal group might involve more than random accumulation through continuous non-zoonotic transmission and may not be indicative of generation distance from the initial zoonotic parent virus. However, the distribution of mutations across the 18 genomes does suggest that MPXV circulating in humans in Nigeria is diverging into subpopulations which may be partially isolated from each other through demographics and dynamics of transmission. This appears to be very different from the international situation, where transmission may currently be concentrated in a more restricted but highly mobile demographic. In addition, the international outbreak appears to involve transmission predominantly via primary rash and this expected to accelerate transmission chains (*10*), which would affect the apparent rate of accumulation of APOBEC3 style mutations. This bears further analysis against genomes from Nigeria concurrent with those from the international expansion and, if available, genomes from human isolates taken in 2020 and 2021. In this context it is interesting that two genomes isolated in the US in 2022 (ON674051 and ON675438; “2022 Other”, in Figure 2 and Tables S2 & S3) from importations unrelated to the wider global outbreak and now designated Lineage A.2 (*12*), have a lower frequency of APOBEC3 style mutations than B.1 isolates.

**Fig. 2.**
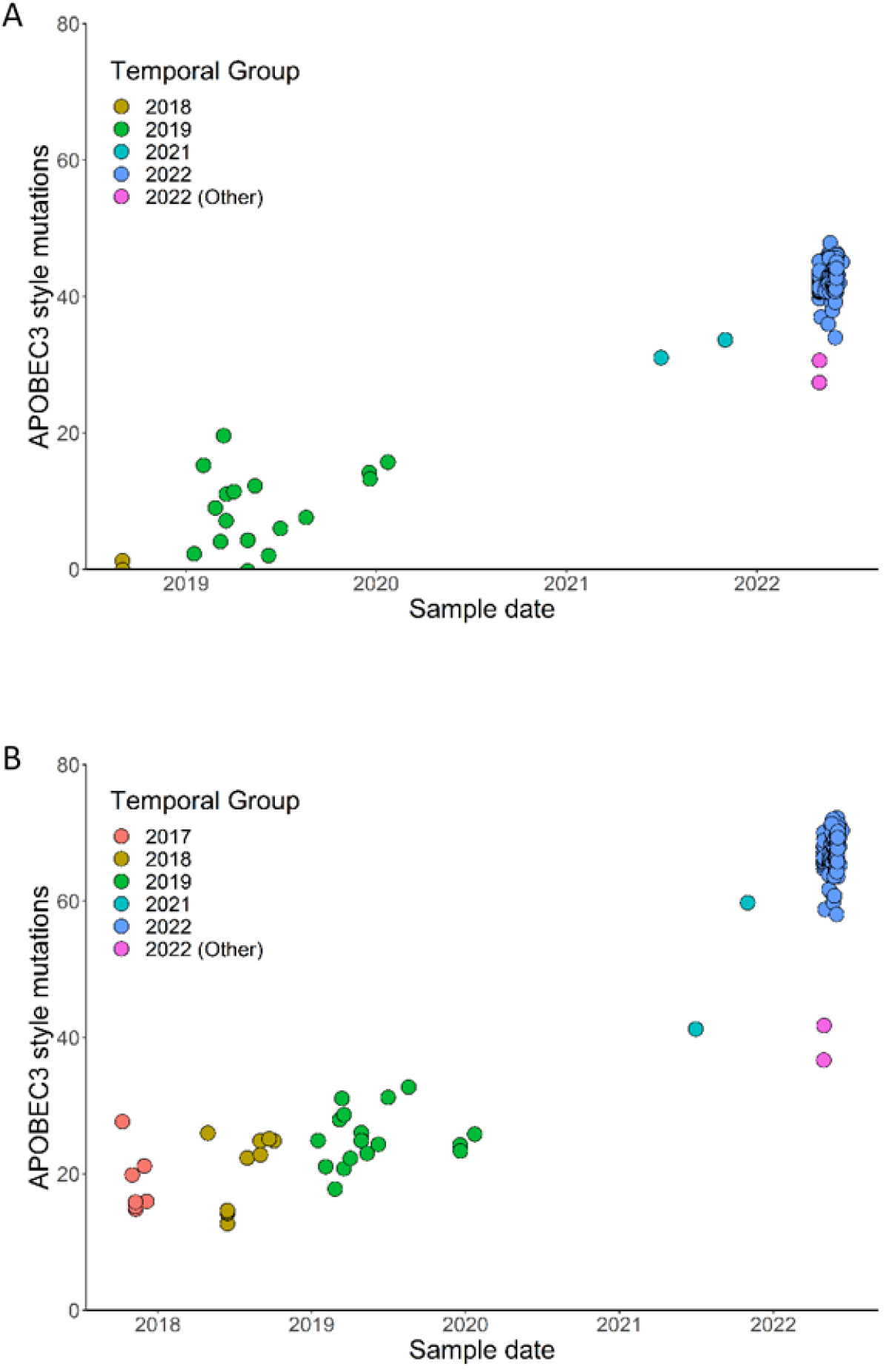
Frequency of APOBEC3 style mutations in clade II genomes isolated since 2019 using the UK 2018 MT903345 isolate as a baseline (A), and in clade II genomes isolated since 2017 using the Nigeria 1971 KJ642617 as a baseline. Genomes are placed in Temporal groups by year of isolation with the exception of OP612691 which is included in the 2019 group but was isolated in January 2020. “Other” refers to Lineage A.2 genomes isolated in 2022 in the US which are phylogenetically separated from the majority of sequences in the international outbreak and arise from a separate lineage within clade II.

**Fig. 3.**
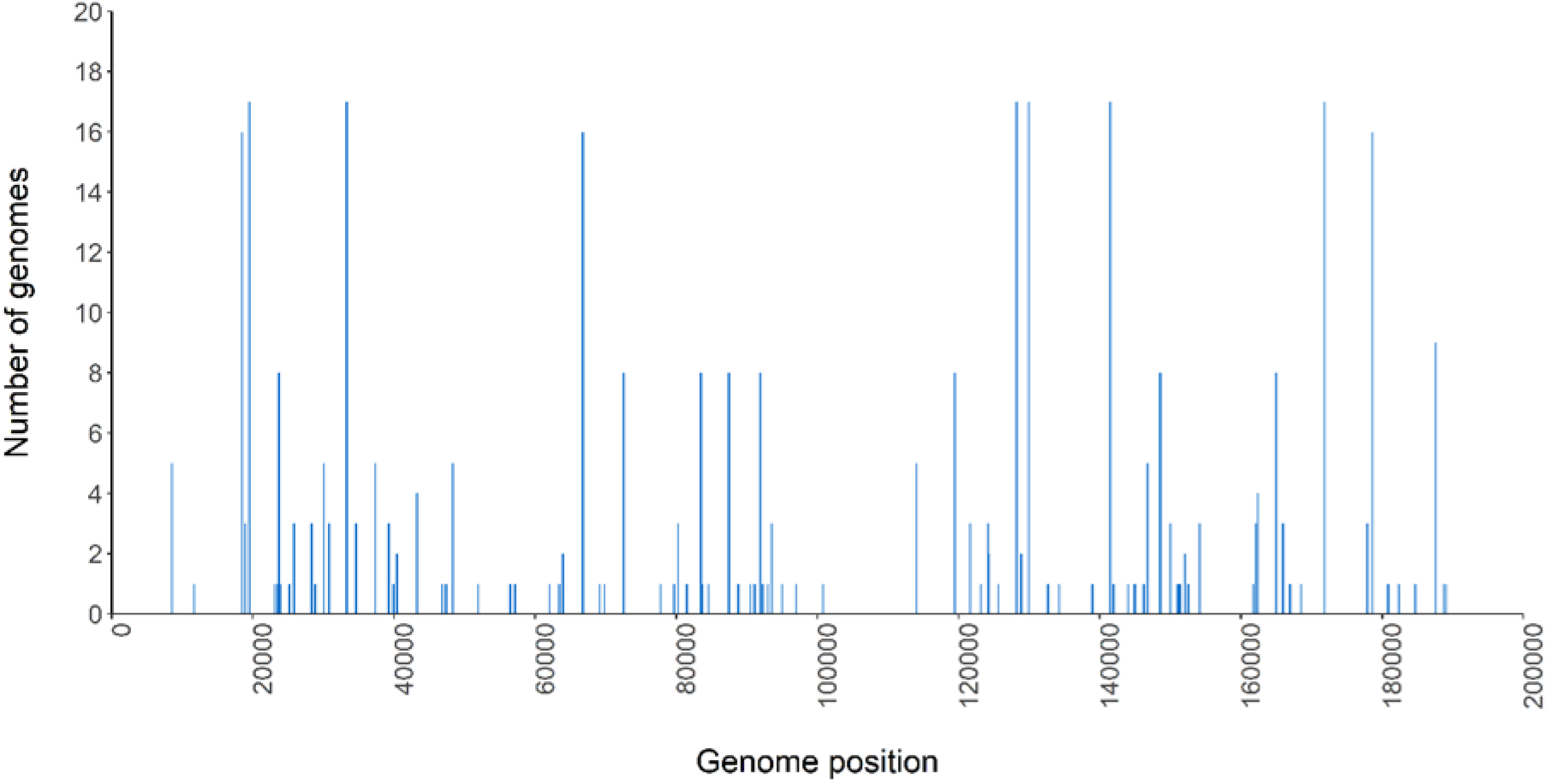
Distribution of APOBEC3 style mutations across the genome. The linear position of all APOBEC3 style mutations for all 18 isolates described in this study is plotted, with the number of isolates with a given mutation shown on the vertical axis. Genome length is 195 Kb.

The possibility that APOBEC3 enzymes may be active on MPXV in humans is an important consideration for our understanding of the potential for MPXV to adapt in humans. Poxvirus genomes are replicated by a virus-encoded proof-reading DNA polymerase, and this is expected to limit the ability of the virus to adapt after a species jump. Mounting evidence that APOBEC3 enzymes may be active on the MPXV genome in human infections indicates an unexpected source of variation that provides raw material for evolution and adaptation independent of the proof-reading polymerase.

### Lineage A.2 in West Africa: A further circulating variant of MPXV

A recent study of MPXV genomes from patients in the US prior to and concurrent with the global outbreak but directly imported from West Africa, has demonstrated that multiple Clade II viruses have been found outside Africa since 2017, independent of the global B.1 lineage (*12*). In particular, three Clade IIb viruses from the US cluster closely in a phylogenetic tree and are designated as Lineage A.2. The history of these three importations indicates they were independently exported from Nigeria over a period of 10 months, suggesting this lineage was circulating in Nigeria prior to the global B.1 outbreak, and it is of note that one of the importations in May 2022 was a female patient recently returned from Nigeria (*12*).

Three of the genomes in this study cluster on the same branch of the phylogenetic tree as the US A.2 genomes and are thus part of this lineage (Figure 1). Importantly, these three Nigerian genomes were isolated between December 2019 and January 2020. This makes it clear that lineage A.2 originated in West Africa and extends the period for which the lineage is known to have circulated in humans to 30 months, during which it has achieved multiple exportations to North America.

Analysis of the six genomes now available for lineage A.2 shows 59 SNPs that are unique to one of the six in comparison with the others, of which 54 (91.5%) are consistent with APOBEC3 action. Five of the six possess at least one SNP not found in any of the others, with OP612690 (67 SNPs relative to KJ642617) having only SNPs that are common to other members.

OP612689 and OP612690 have one and two unique SNPs respectively, while the later three genomes from July 2021 and May 2022 in the USA have 21, 19, and 19 SNPs that are unique to them relative to the other genomes in the lineage. Although there are only six genomes for comparison, the distribution of mutations between them indicate that two of the three Lineage A.2 genomes presented in this study are not directly ancestral to the three later A.2 genomes, and the six viruses are members of a lineage that is further diversifying.

To investigate these viruses further we undertook a more detailed analysis, using only concatenated protein amino acid sequences from complete genomes, and excluding genomes with long runs of ambiguous nucleotides in the assembled sequences. This provides a more granular picture than nucleotide-based trees, although the number of genomes for comparison is reduced. With this approach these six viruses form a cluster on a long branch separating them from other clade IIb.A viruses (Figure 4), supporting the lineage A.2 designation given to the three USA isolates in the cluster by Gigante *et al*. (*12*) In addition, using this analysis two further clusters appear to have branched away and may potentially be regarded as distinct lineages. One of these groups 10 of the viruses in this study, and the other groups two of the viruses in this study along with 2018 isolates from Israel, Nigeria and the UK. Although these two clusters do not have an identifying characteristic like the A.2 lineage (see below), taken with the A.2 and B.1 lineages they are a clear indication that multiple lineages of MPXV have achieved sustained human-to-human transmission from the presumed single zoonotic event, and have co-existed in humans over a period of several years. This suggests the global success of the B.1 lineage over other co-existing lineages may be due at least in part to a founder effect from chance introduction to the international MSM community.

**Fig. 4.**
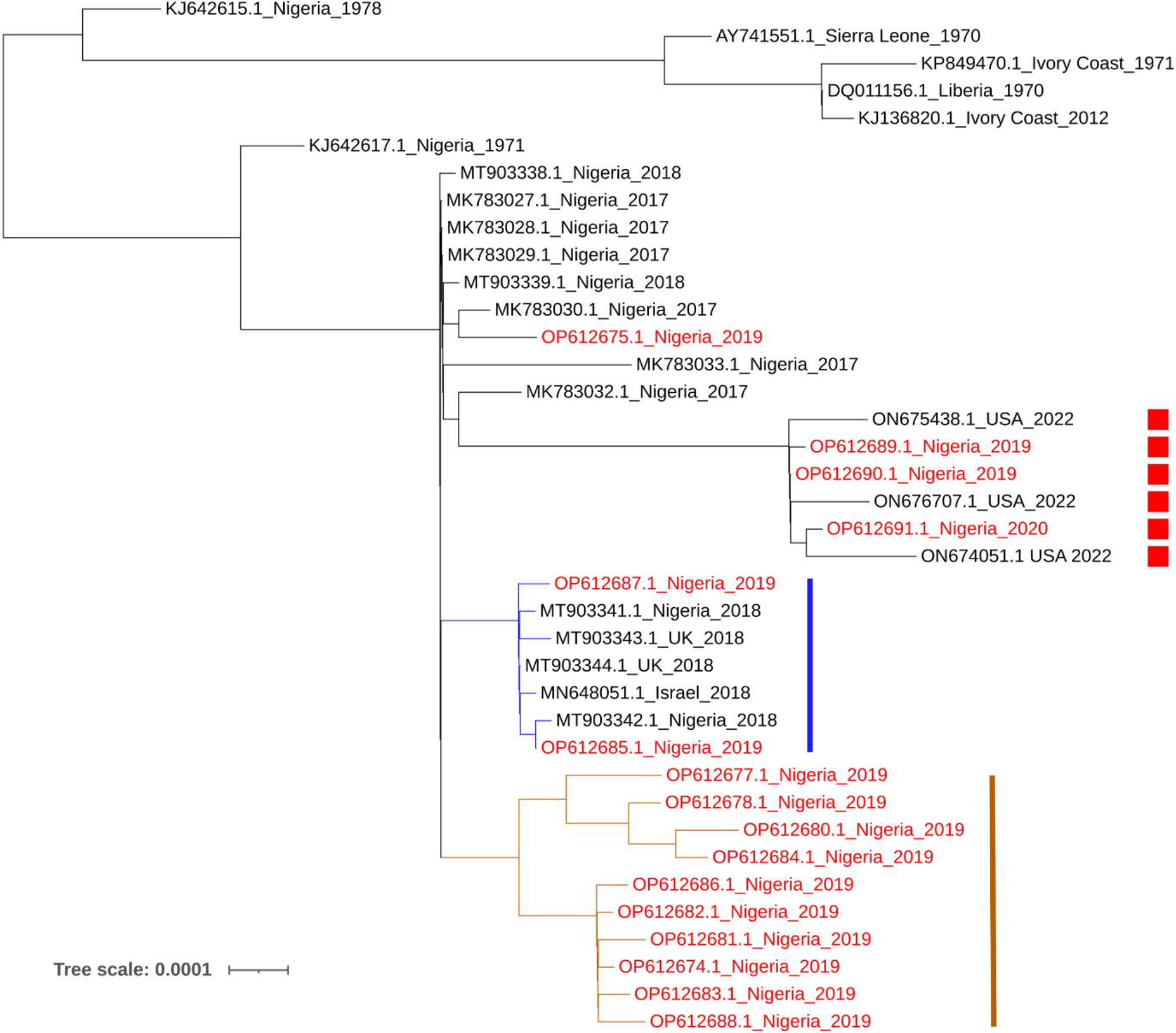
Phylogenetic Tree of Lineage A MPXV genomes. The tree is generated using concatenated protein amino acid sequences from complete genomes with a limited number of unknown bases in their sequence, and excluding genes with any ambiguous nucleotides in the assembled genome. Genomes described in this study are highlighted in red and those with a disrupted A46R gene (Lineage A.2) are marked with a red square. Two additional potential lineages are marked with a blue or brown bar.

### Gene disruption in lineage A2

The three Lineage A.2 genomes presented here have a nonsense mutation in the A46R gene (Copenhagen nomenclature (*20*)). This gene is highly conserved between MPXV and Vaccinia virus (VACV), and encodes a TOLL/IL1-receptor (TIR) homologue. The A46 protein interferes with IL1-depedent signal transduction and activation of NF-KB and type-I interferon (IFN-I) responses, and deletion mutants in VACV are attenuated for virulence in mice (*21, 22*). A46 is disrupted in three of the 18 isolates (OP612690, OP612689 and OP612691), with an identical nonsense SNP that is compatible with APOBEC3 activity (Table S1). All were collected between December 2019 and January 2020. OP612690 and OP612689 were from individuals in Imo and Rivers State respectively, and are adjacent on the phylogenetic tree. Geographic metadata is missing for OP612691, but the genome is in the same phylogenetic cluster with OP612690 and OP612689. Importantly, an identical mutation is seen in the three the A.2 isolates from the United States ON676707 (July 2021), ON674051 and ON675438 (May 2022), and these three, which are independent of the B.1 lineage (*23*), form a contiguous cluster with OP612690, OP612689 and OP612691 on the phylogenetic tree (Figures 1A and 4).

APOBEC3 style mutations described so far in the B.1 lineage occur at TC dinucleotides, with the C nucleotide deaminated to T (*11, 17*). TC dinucleotides are common in the MPXV genome. The frequency of TC to TT changes, and the increase over time, suggest the process is relatively inefficient, with only a small number of mutations arising in a replication cycle. The distribution of APOBEC3 style mutations in the various genomes reported here and elsewhere suggest the specific TC dinucleotides mutated to TT in a replication cycle is effectively random, and the radiation of MPXV in human populations will result in continuous and measurable divergence with respect to this type of mutation. The presence of an identical APOBEC3 style mutation in all six Lineage A.2 genomes suggests it is not independent, and coupled with their isolation over a period from December 2019 to May 2022, that this A46 disruption variant is actively circulating between human hosts. For many SNPs, this might mean they are neutral with respect to adaptation. However a mutation that disrupts an immune modulator protein is not expected to be neutral *in vivo*, and there is thus a possibility that the disruption of A46R has adaptive value for the virus.

MPXV encodes a plethora of proteins that have been shown in other OPVs to modulate innate immunity, including several that interfere with IL1 responses. In experimental systems, disrupting these genes is often attenuating in laboratory mice. However disruptions are also seen in natural systems, most notably in *Variola virus* (VARV), the causative agent of smallpox, where sequencing studies have shown that several such genes are disrupted, including genes that engage with IL1 (*24, 25*). The adaptive value of these gene disruptions in natural systems is not clear. Although only 3/18 sequences in this study and three from elsewhere have this disruption, it is nevertheless noteworthy that it is an APOBEC3 style nonsense mutation in six phylogenetically close genomes, representing a variant which has survived in human transmission chains for more than two years. It is known that OPVs adapt to new hosts through a process of gene loss, analogous to the disruption of A46R (*26*). Consequently, sustained transmission of an A46R disruption variant may represent a step on the path to adaption, and the possibility of B.1 viruses in the global outbreak acquiring such mutations, or other lineages that have already acquired gene disruptions acquiring additional gene disruptions, is a cause for concern.

The dynamics with which variants emerge and displace parents in a natural system are not understood for poxviruses, but viruses transmitted by sustained skin-to-skin contact will have different transmission dynamics to respiratory infections such as SARS-COV2, and this is likely to delay the spread of variants. It is possible that in West Africa, with relatively low rates of human-to-human transmission, favourable variants are lost by chance, or have a protracted lag in establishment. If variants with mutations analogous to disruption of A46R are mildly adaptive for non-zoonotic human-to-human transmission, their apparent low prevalence may be due to founder effects facilitating other variants by chance.

In the last century human MPXV infection is assumed to have been suppressed by the circulation of smallpox, and subsequently by the artificial “circulation” of smallpox vaccination. With smallpox eradication and the cessation of vaccination, the human niche has been open, and as anti-smallpox immunity has waned, the incidence of Mpox zoonosis has steadily increased (*27*). Although human infection has been rare in West Africa, the situation has changed with the sudden and continuing spike of human infections in Nigeria in 2017, which is now expanding globally. Our findings demonstrate that as of 2019 multiple variants of MPXV were circulating in West Africa with a level of APOBEC-like mutations consistent with sustained non-zoonotic transmission. The inactivation of an immunomodulating gene in Lineage A.2 viruses separated by 30 months over two continents suggests continuing circulation, and active selection pressure to refine fitness and spread between humans without reintroduction from reservoirs. To date, variants of the type described here are not present in lineages descending from the 2022 lineage B.1 outbreak that has spread globally. However, we should be aware of the possibility of their genesis, especially if sexual contact plays a greater role in the global expansion success of B.1 than in the sustained human transmission observed in West Africa.

In conclusion, our understanding of the evolution of MPXV during sustained human-to-human transmission is hampered by the lack of genome sequences from 2019 to 2021, and this is ameliorated by 18 genomes from January 2019 to January 2020 provided here, which may the only genomes available from this period. These confirm the incidence of mutations consistent with APOBEC3 activity in all phases of the outbreak post 2017 for which genomes are available; they suggest the outbreak may stem from a single zoonotic event, with subsequent splits into partially isolated virus populations in Nigeria that may be evolving independently of each other and diverging. The prevalence of APOBEC3 style mutations also suggests a mutational mechanism that compensates for the stability anticipated from a proof-reading polymerase. In addition, disruption of an immune modulating gene, a process shown to have occurred in the adaptation of OPVs to new hosts, has been observed in a cluster of six genomes; this represents an additional lineage that has survived for more than two years in West Africa and has been exported to the North America three times, including via a female patient. The variation among isolates, and the long-term survival of an A46R disruption variant, suggest that a chance founder effect may be a large part of the success of lineage B.1 over other lineages in the global expansion of MPXV in the MSM community. It is anticipated that if Lineage B.1 is eliminated, Lineage A MPXV will continue to evolve if its human transmission chain is not broken, and there is no *a priori* reason to expect transmission of future variants will be limited to close or extended skin-to-skin contact.

## Acknowledgments

We would like to thank Baharak Afshar for expert help with liaison arrangements and shipping between NCDC and UKHSA.

## Author contributions

Sample collection: NN, AYO, CC, AA, OB, NM, OB, OA Sequencing: NN, KL, CC, AA, OB, NM, OB, OA Bioinformatics and Phylogenetics: KL, DK, PR, RPS, AR Conceptualization: NN, JA, AYO, PR, CMM, BG, CI, IA, DOU Methodology: NN, JA, KL, AYO, AA, DK, PR, TM, Statistical Analysis: DK, TM Funding acquisition: JA, STP, BG, CB, CI Project administration and leadership: NN, JA, AA, BG, CI, DOU Writing – original draft: DK, CMM, PR, DOU Writing – review & editing: NN, JA, KL, AYO, DK, CMM, PR, RPS, TM, CI, IA, DOU Crown Copyright ©2022

## Data and materials availability

Genomes for the 18 isolates described here are deposited in GenBank; accession numbers are listed in Table S5.

## Supplementary Materials

Materials and Methods

Tables S1 to S5

## References and Notes

1. Z. Jezek, F. Fenner, Human monkeypox. (Karger Publishers, 1988).

2. D. Ogoina et al., Clinical course and outcome of human monkeypox in Nigeria. Clinical Infectious Diseases 71, e210–e214 (2020).

3. A. Yinka-Ogunleye et al., Outbreak of human monkeypox in Nigeria in 2017–18: a clinical and epidemiological report. The Lancet Infectious Diseases 19, 872–879 (2019).

4. M. R. Mauldin et al., Exportation of monkeypox virus from the African continent. The Journal of infectious diseases 225, 1367–1376 (2022).

5. O. T. Ng et al., A case of imported Monkeypox in Singapore. The Lancet Infectious Diseases 19, 1166 (2019).

6. A. Vaughan et al., Two cases of monkeypox imported to the United Kingdom, September 2018. Eurosurveillance 23, 1800509 (2018).

7. R. Vivancos et al., Community transmission of monkeypox in the United Kingdom, April to May 2022. Eurosurveillance 27, 2200422 (2022).

8. N. Erez et al., Diagnosis of imported monkeypox, Israel, 2018. Emerging infectious diseases 25, 980 (2019).

9. M. P. Duque et al., Ongoing monkeypox virus outbreak, Portugal, 29 April to 23 May 2022. Eurosurveillance 27, 2200424 (2022).

10. D. O. Ulaeto, J. Dunning, M. W. Carroll, Evolutionary implications of human transmission of monkeypox: the importance of sequencing multiple lesions. The Lancet Microbe, (2022).

11. J. Isidro et al., Phylogenomic characterization and signs of microevolution in the 2022 multi-country outbreak of monkeypox virus. Nature Medicine, 1–1 (2022).

12. C. M. Gigante et al., Multiple lineages of monkeypox virus detected in the United States, 2021–2022. Science, eadd4153 (2022).

13. C. Happi et al., Urgent need for a non-discriminatory and non-stigmatizing nomenclature for monkeypox virus. Virological. org, (2022).

14. L. A. S. S. O. Va, Frace AM Li Y 2005 A tale of two clades: monkeypox viruses. J Gen Virol 86, 26612672.

15. S. O. Foster et al., Human monkeypox. Bulletin of the World Health Organization 46, 569 (1972).

16. Y. Nakazawa et al., A phylogeographic investigation of African monkeypox. Viruses 7, 2168–2184 (2015).

17. A. O’Toole, A. Rambaut, in https://scanmail.trustwave.com/?c=7369&d=qNjW4kQovAKJ5KdfgmUZVoNWASfZiuixFJgB2qNMRg&u=https%3a%2f%2fvirological%2eorg%2ft%2f830. (Virological.org, 2022).

18. A. O’Toole, A. Rambaut, in https://virological.org/t/an-apobec3-molecular-clock-to-estimate-the-date-of-emergence-of-hmpxv/885. (Virological.org, 2022).

19. I. Cohen-Gihon et al., Identification and whole-genome sequencing of a Monkeypox virus strain isolated in Israel. Microbiology Resource Announcements 9, e01524–01519 (2020).

20. S. J. Goebel et al., The complete DNA sequence of vaccinia virus. Virology 179, 247–266 (1990).

21. J. Stack et al., Vaccinia virus protein A46R targets multiple Toll-like–interleukin-1 receptor adaptors and contributes to virulence. The Journal of experimental medicine 201, 1007–1018 (2005).

22. A. Bowie et al., A46R and A52R from vaccinia virus are antagonists of host IL-1 and toll-like receptor signaling. Proceedings of the National Academy of Sciences 97, 10162–10167 (2000).

23. C. M. Gigante et al., Multiple lineages of Monkeypox virus detected in the United States, 2021-2022. bioRxiv, (2022).

24. A. Alcami, G. L. Smith, Comment on the paper by Shchelkunov et al.(1993) FEBS Letters 319, 80-83. Two genes encoding poxvirus cytokine receptors are disrupted or deleted in variola virus. FEBS letters 335, 136–137 (1993).

25. B. Mühlemann et al., Diverse variola virus (smallpox) strains were widespread in northern Europe in the Viking Age. Science 369, eaaw8977 (2020).

26. R. C. Hendrickson, C. Wang, E. L. Hatcher, E. J. Lefkowitz, Orthopoxvirus genome evolution: the role of gene loss. Viruses 2, 1933–1967 (2010).

27. A. W. Rimoin et al., Major increase in human monkeypox incidence 30 years after smallpox vaccination campaigns cease in the Democratic Republic of Congo. Proceedings of the National Academy of Sciences 107, 16262–16267 (2010).

28. T. Charalampous et al., Nanopore metagenomics enables rapid clinical diagnosis of bacterial lower respiratory infection. Nature biotechnology 37, 783–792 (2019).

29. A. L. Greninger et al., Rapid metagenomic identification of viral pathogens in clinical samples by real-time nanopore sequencing analysis. Genome medicine 7, 1–13 (2015).

30. L. E. Kafetzopoulou et al., Assessment of metagenomic Nanopore and Illumina sequencing for recovering whole genome sequences of chikungunya and dengue viruses directly from clinical samples. Eurosurveillance 23, 1800228 (2018).

31. M. S. Wright et al., SISPA-Seq for rapid whole genome surveys of bacterial isolates. Infection, Genetics and Evolution 32, 191–198 (2015).

32. J. Quick. (2018).

33. S. Koren et al., Canu: scalable and accurate long-read assembly via adaptive k-mer weighting and repeat separation. Genome research 27, 722–736 (2017).

34. S. F. Altschul, W. Gish, W. Miller, E. W. Myers, D. J. Lipman, Basic local alignment search tool. Journal of molecular biology 215, 403–410 (1990).

35. A. R. Quinlan, I. M. Hall, BEDTools: a flexible suite of utilities for comparing genomic features. Bioinformatics 26, 841–842 (2010).

36. K. Katoh, D. M. Standley, MAFFT multiple sequence alignment software version 7: improvements in performance and usability. Molecular biology and evolution 30, 772–780 (2013).

37. L.-T. Nguyen, H. A. Schmidt, A. Von Haeseler, B. Q. Minh, IQ-TREE: a fast and effective stochastic algorithm for estimating maximum-likelihood phylogenies. Molecular biology and evolution 32, 268–274 (2015).

38. A. Stamatakis, RAxML version 8: a tool for phylogenetic analysis and post-analysis of large phylogenies. Bioinformatics 30, 1312–1313 (2014).

39. R. C. Team. (2019).

40. D. Ogle, P. Wheeler, A. Dinno. (Vienna: R Core Team., 2022).

