## Supplementary material for "Different Coexisting Mpox Lineages Were Continuously Circulating in Humans Prior to 2022": MM and Supplementary Tables S1-S5

**The PDF file includes:**

Materials and Methods
Tables S1 to S5

### Materials and Methods

Sample collection and Nucleic acid extraction

Twenty lesion swab samples were selected for sequencing. Samples were selected based on sample CT on PCR and collection date to give as wide a temporal range as possible at the time. A CT of between 14.90 (OP612675) and 19.98 (OP612677) (mean: 17.82) was recorded for samples. Prior to extraction, background host DNA was removed with an adapted nuclease treatment (*28*), incubating with HL-SAN Nuclease (ArticZymes) at a final concentration of 1U/ul, for 15 minutes at 37°C. Samples were then inactivated and extracted using QIAamp DNA Mini Kit (Qiagen), as per manufacturer's instructions.

Sequencing

In order to produce sufficient quantities of material to sequence, DNA samples were tagged, randomly primed and amplified using a modified sequence-independent amplification (*29-31*). A total of five sequencing runs using the Oxford Nanopore Technologies (ONT™) MinION sequencer and R9.41 flowcells were carried out, with each containing five Mpox samples. Sequencing libraries for each were prepared using the ONT Native barcoding expansion (EXP-NDB103) with the ligation sequencing kit (SQK-LSK109) and a modified One-pot protocol (*32*).

Bioinformatics

Fastq reads were basecalled and de-multiplexed using guppy (version 3.6) (ONT), with reads under a qScore of q7 being discarded. Following this, reads equal and above 1000bp were assembled using Canu (version 2.1.1) (*33*) with generated Contigs being analysed in BLASTn (*34*) to select a suitable Mpox reference sequence for sequence read alignment. For cases where Canu failed to produce any Mpox Contigs, the NCBI RefSeq MPXV sequence (NC_003310) was used as a reference

The Medaka (version 1.2.2) program (available from https://github.com/nanoporetech/medaka ), using the medaka consensus command and the r941_min_high_g360 model, was used to generate a polished consensus sequence. Multiple rounds of polishing were performed to improve the consensus and close gaps while incorporating INDELs. Following the final round of Medaka polishing, BEDtools (version 2.27.1) (*35*) was used ascertain coverage across the sequenced genome. Bases with a read depth of <10 were regarded as indeterminate and replaced with "N" using an in-house R-script.

Following ONT MinION sequencing, between $5.63 \times 10^4$ (OP612687) and $3.60 \times 10^6$ (OP612691) fastq reads (mean: $1.36 \times 10^6$) above a qScore of q7 were generated. Following alignment to a respective reference sequence, low sequence coverage was identified for two samples (MPXV/100/19 (0.1%) and MPXV/050/19) (50.20%). These samples were removed from all downstream analysis. For the remaining 18 samples, between 90.76% (OP612676) and 100% (OP612690 & OP612691) (mean: 99.19%) of the genome was generated with an average sequence read coverage of 1475).

Sequences were aligned with 337 previously sequenced Mpox isolates using MAFFT (*36*) (version 7.453) (maximum iterations = 15) before the first 14,619bp and last 17,148bp of all sequences were trimmed. Following this, a maximum likelihood phylogenetic tree showing the evolutionary relationship between aligned MPXV isolates was generated using IQ-Tree (*37*) (version 2.1.3) (MFP model selected, replicates for bootstrap: 1000). Finally, all output genomes were compared against a sequence characterised Clade II isolate (KJ642617) to characterise the SNP and INDEL differences. KJ642617 is from the Clade II, isolated in from a human case in Nigeria in 1971.

We also generated a whole-genome tree based on coding regions by identifying a smaller set of high-quality genomes with a limited number of unknown bases in their sequence. We reannotated them with an in-house script, using the annotation of ON954773 as a starting point. Protein sequences still present in all genomes after removal of ambiguous/unknown amino acids (OPG024, OPG027, OPG029, OPG030, OPG031, OPG034, OPG035, OPG036, OPG037, OPG038, OPG039, OPG040, OPG042, OPG043, OPG044, OPG045, OPG046, OPG047, OPG048, OPG049, OPG050, OPG051, OPG052, OPG053, OPG054, OPG055, OPG056, OPG057, OPG058, OPG059, OPG060, OPG061, OPG062, OPG065, OPG066, OPG069, OPG070, OPG073, OPG075, OPG076, OPG077, OPG079, OPG081, OPG082, OPG083, OPG085, OPG086, OPG087, OPG089, OPG090, OPG091, OPG092, OPG093, OPG094, OPG095, OPG096, OPG097, OPG098, OPG099, OPG101, OPG102, OPG104, OPG105, OPG106, OPG107, OPG108, OPG109, OPG110, OPG111, OPG112, OPG113, OPG114, OPG115, OPG116, OPG118, OPG119, OPG120, OPG121, OPG122, OPG123, OPG124, OPG125, OPG126, OPG127, OPG128, OPG129, OPG130, OPG131, OPG132, OPG133, OPG134, OPG137, OPG138, OPG139, OPG140, OPG141, OPG142, OPG143, OPG144, OPG146, OPG147, OPG148, OPG149, OPG150, OPG151, OPG154, OPG155, OPG156, OPG157, OPG158, OPG159, OPG160, OPG161, OPG162, OPG163, OPG164, OPG165, OPG166, OPG167, OPG170, OPG171, OPG173, OPG174, OPG175, OPG176, OPG178, OPG180, OPG181, OPG187, OPG189, OPG190, OPG191, OPG192, OPG193, OPG195, OPG198, OPG199, OPG200, OPG205, OPG209) were collected and concatenated for each genome. The concatenated sequences were aligned with MAFFT (*36*) and, from the result, a maximum likelihood phylogenetic tree was produced with RAxML (*38*) version 8 using a protein GTR+gamma model.

To characterise any APOPBEC3 style mutations (TC > TT and GA > AA) which have recently evolved, all generated NCDC and 2022 outbreak samples were aligned to the UK_P1, UK_P2 and UK_P3 (MT903343, MT903344, MT903345) MPXV genomes, sequenced from samples collected in the UK in 2018, and separately against KJ642617. A sliding window analysis (window size: 2, step size: 1) was performed using an in-house R-script to identify any associated APOBEC3 style dinucleotide changes.

<u>Additional Genomes</u>

All genomes used in this analysis are listed in Supplementary Table 5. Genomes used for comparison with the 18 isolates described here were downloaded from GenBank, including ON676707, ON674051 and ON675438 (*23*). KJ642617 is used for comparisons because it is the earliest available sequence from Nigeria, and is assumed to be directly zoonotic and thus not multiply passaged between humans.

<u>Statistical Analysis</u>

Summary statistics on APOBEC3 style mutations were presented by temporal group. As APOBEC3 style mutations were not normally distributed the median, range and inter-quartile range (IQR) were presented. A Kruskal-Wallis test was first used to determine if there were significant differences between temporal groups, followed by pairwise comparisons between temporal groups made using Dunn's tests, with a Bonferroni correction applied to adjust for multiple testing. Statistical analysis was carried out in R (*39*) using the kruskal.test function from the R base package, and the DunnTest function from the FSA package (*40*).

### Table S1.

| Genome position | Sample | Type | Reference sequence | Alternate sequence | Effect | Gene product |
|---|---|---|---|---|---|---|
| 6744 | OP612679 | del | CT | C | | |
| 7110 | OP612688 | snp; APOBEC3 (possible error) | C | T | | |
| 8547 | OP612675, OP612676, OP612678, OP612680, OP612684 | snp; APOBEC3 | C | T | | |
| 8662 | OP612679 | snp; APOBEC3 (possible error) | C | T | | |
| 9055 | OP612674, OP612675, OP612676, OP612677, OP612678, OP612680, OP612681, OP612682, OP612683, OP612684, OP612685, OP612686, OP612687, OP612688, OP612689, OP612690, OP612691 | snp | C | A | | |
| 11717 | OP612674 | snp; APOBEC3 | C | T | R449Q | ankyrin-like protein |
| 13460 | OP612689, OP612690, OP612691 | snp | C | T | | |
| 13787 | OP612674, OP612675, OP612676, OP612677, OP612678, OP612680, OP612681, OP612682, OP612683, OP612684, OP612685, OP612686, OP612687, OP612688, OP612689, OP612690, OP612691 | snp | A | G | M548T | ankyrin-like protein |
| 16770 | OP612674, OP612675, OP612676, OP612677, OP612678, OP612680, OP612681, OP612682, OP612683, OP612684, OP612685, OP612686, OP612687, OP612688, OP612689, OP612690, OP612691 | del | GTCATCA | G | D155_D156(del) | hypothetical protein |

| | | | | | | |
|---|---|---|---|---|---|---|
| 18480 | OP612674, OP612675, OP612677, OP612678, OP612679, OP612681, OP612682, OP612683, OP612684, OP612685, OP612686, OP612687, OP612688, OP612689, OP612690, OP612691 | snp; APOBEC3 | G | A | L173L | hypothetical protein |
| 18960 | OP612689, OP612690, OP612691 | snp; APOBEC3 | C | T | D13N | hypothetical protein |
| 19535 | OP612674, OP612675, OP612676, OP612677, OP612678, OP612680, OP612681, OP612682, OP612683, OP612684, OP612685, OP612686, OP612687, OP612688, OP612689, OP612690, OP612691 | snp; APOBEC3 | C | T | S213S | hypothetical protein |
| 23139 | OP612681 | snp; APOBEC3 | G | A | I63I | hypothetical protein |
| 23521 | OP612677 | snp; APOBEC3 | C | T | D256N | ankyrin-like protein |
| 23732 | OP612674, OP612679, OP612681, OP612682, OP612683, OP612686, OP612687, OP612688 | snp; APOBEC3 | C | T | S185S | ankyrin-like protein |
| 23879 | OP612681 | snp; APOBEC3 | G | A | V136V | ankyrin-like protein |
| 25240 | OP612691 | snp; APOBEC3 | C | T | R136Q | serine protease inhibitor-like protein SPI-3 |
| 25829 | OP612679, OP612686, OP612687 | snp; APOBEC3 | G | A | | |
| 28332 | OP612675, OP612676, OP612678 | snp; APOBEC3 | C | T | S25F | B15R-like protein |
| 28813 | OP612674 | snp; APOBEC3 | G | A | I206I | hypothetical protein |
| 29916 | OP612674, OP612675, OP612676, OP612677, OP612678, OP612680, OP612681, OP612682, OP612683, OP612684, OP612685, OP612686, OP612688, OP612689, OP612690, OP612691 | del | ACCATT | A | N483fs | kelch-like protein |

| | | | | | | |
|---|---|---|---|---|---|---|
| 30062 | OP612675, OP612676, OP612678, OP612680, OP612684 | snp; APOBEC3 | G | A | P436S | kelch-like protein |
| 30809 | OP612689, OP612690, OP612691 | snp; APOBEC3 | C | T | D187N | kelch-like protein |
| 32381 | OP612674, OP612675, OP612676, OP612677, OP612678, OP612680, OP612681, OP612682, OP612683, OP612684, OP612685, OP612686, OP612687, OP612688, OP612689, OP612690, OP612691 | ins | T | TGAC | S313_H314(ins)R | hypothetical protein |
| 33313 | OP612674, OP612675, OP612676, OP612677, OP612678, OP612680, OP612681, OP612682, OP612683, OP612684, OP612685, OP612686, OP612687, OP612688, OP612689, OP612690, OP612691 | snp; APOBEC3 | C | T | T3T | hypothetical protein |
| 33347 | OP612681 | snp; APOBEC3 | G | A | | |
| 34642 | OP612689, OP612690, OP612691 | snp; APOBEC3 | C | T | V73V | hypothetical protein |
| 34748 | OP612689, OP612690, OP612691 | snp; APOBEC3 | C | T | G38E | hypothetical protein |
| 37379 | OP612675, OP612676, OP612678, OP612680, OP612684 | snp; APOBEC3 | G | A | S609F | hypothetical protein |
| 38237 | OP612674, OP612675, OP612676, OP612677, OP612678, OP612680, OP612681, OP612682, OP612683, OP612684, OP612685, OP612686, OP612687, OP612688, OP612689, OP612690, OP612691 | snp | G | A | A323V | hypothetical protein |
| 39289 | OP612679, OP612686, OP612687 | snp; APOBEC3 | C | T | E359E | palmytilated EEV membrane protein |
| 39937 | OP612685 | snp; APOBEC3 | C | T | T143T | palmytilated EEV membrane protein |

| | | | | | | |
|---|---|---|---|---|---|---|
| 40421 | OP612680, OP612684 | snp; APOBEC3 | C | T | D62N | hypothetical protein |
| 43288 | OP612675, OP612678, OP612680, OP612684 | snp; APOBEC3 | G | A | L191L | poly-A polymerase catalytic subunit VP55 |
| 46855 | OP612678 | snp; APOBEC3 | G | A | F248F | bifunctional DNA-dependent RNA polymerase subunit rpo30 |
| 47065 | OP612674, OP612675, OP612676, OP612677, OP612678, OP612680, OP612681, OP612682, OP612683, OP612684, OP612685, OP612686, OP612687, OP612688, OP612689, OP612690, OP612691 | snp | G | A | D178D | bifunctional DNA-dependent RNA polymerase subunit rpo30 |
| 47394 | OP612688 | snp; APOBEC3 | C | T | E69K | bifunctional DNA-dependent RNA polymerase subunit rpo30 |
| 48308 | OP612674, OP612675, OP612676, OP612677, OP612678, OP612680, OP612681, OP612682, OP612683, OP612684, OP612685, OP612686, OP612687, OP612688, OP612689, OP612690, OP612691 | snp | A | G | | |
| 48350 | OP612675, OP612676, OP612678, OP612680, OP612684 | snp; APOBEC3 | G | A | | |
| 48697 | OP612674, OP612675, OP612676, OP612677, OP612678, OP612680, OP612681, OP612682, OP612683, OP612684, OP612685, OP612686, OP612687, OP612688, OP612689, OP612690, OP612691 | snp | C | A | T109T | hypothetical protein |
| 51979 | OP612677 | snp; APOBEC3 | C | T | R877Q | DNA polymerase |
| 53377 | OP612674, OP612675, OP612676, OP612677, OP612678, OP612680, | snp | C | A | W411L | DNA polymerase |

| | | | | | | |
|---|---|---|---|---|---|---|
| | OP612681, OP612682, OP612683, OP612684, OP612685, OP612686, OP612687, OP612688, OP612689, OP612690, OP612691 | | | | | |
| 56521 | OP612677 | snp; APOBEC3 | C | T | E259K | hypothetical protein |
| 57138 | OP612677 | snp; APOBEC3 | G | A | S53L | hypothetical protein |
| 62014 | OP612677 | snp; APOBEC3 | G | A | L59L | large subunit of ribonucleotide reductase protein |
| 63434 | OP612680 | snp; APOBEC3 | C | T | L63L | hypothetical protein |
| 63971 | OP612680, OP612684 | snp; APOBEC3 | C | T | E306K | viral core Cysteine proteinase |
| 66750 | OP612674, OP612675, OP612676, OP612677, OP612678, OP612681, OP612682, OP612683, OP612684, OP612685, OP612686, OP612687, OP612688, OP612689, OP612690, OP612691 | snp; APOBEC3 | G | A | R620Q | bifunctional DNA/RNA-helicase/DExH-NPH-II |
| 69187 | OP612682 | snp; APOBEC3 | C | T | S54F | late transcription elongation factor |
| 69848 | OP612677 | snp; APOBEC3 | G | A | F62F | hypothetical protein |
| 72532 | OP612674, OP612679, OP612681, OP612682, OP612683, OP612686, OP612687, OP612688 | snp; APOBEC3 | C | T | D196N | virion structural protein |
| 74202 | OP612677 | snp | G | A | A85T | myristylprotein |
| 76137 | OP612674 | snp | G | A | S303S | hypothetical protein |
| 76248 | OP612674, OP612675, OP612677, OP612678, OP612680, OP612682, OP612683, OP612684, OP612686, OP612687, OP612688, OP612690, OP612691 | Del (possible error) | CT | C | K266fs | hypothetical protein |
| 77805 | OP612681 | snp; APOBEC3 | G | A | D246N | core protein vp8 |
| 77896 | OP612688 | snp; APOBEC3 (possible error) | C | T | S21F | putative membrane protein |

| | | | | | | |
|---|---|---|---|---|---|---|
| 79708 | OP612679 | snp; APOBEC3 | G | A | D152N | bifunctional subunit of multifunctional poly-A polymerase |
| 80269 | OP612689, OP612690, OP612691 | snp; APOBEC3 | G | A | L33L | DNA-dependent RNA polymerase subunit rpo22 |
| 81156 | OP612674, OP612675, OP612676, OP612677, OP612678, OP612680, OP612681, OP612682, OP612683, OP612684, OP612685, OP612686, OP612687, OP612688, OP612689, OP612690, OP612691 | snp | C | T | A12T | late 16 kDa putative membrane protein |
| 81445 | OP612686 | snp; APOBEC3 | G | A | K50K | DNA-dependent RNA polymerase subunit rpo147 |
| 81524 | OP612679 | snp; APOBEC3 | G | A | E77K | DNA-dependent RNA polymerase subunit rpo147 |
| 81693 | OP612683 | snp | G | A | S133N | DNA-dependent RNA polymerase subunit rpo147 |
| 83496 | OP612674, OP612679, OP612681, OP612682, OP612683, OP612686, OP612687, OP612688 | snp; APOBEC3 | C | T | S734L | DNA-dependent RNA polymerase subunit rpo147 |
| 83666 | OP612681 | snp; APOBEC3 | C | T | L791L | DNA-dependent RNA polymerase subunit rpo147 |
| 84178 | OP612677 | snp | C | T | F961F | DNA-dependent RNA polymerase subunit rpo147 |
| 84586 | OP612679 | snp; APOBEC3 | G | A | K1097K | DNA-dependent RNA polymerase subunit rpo147 |
| 87197 | OP612675, OP612676, OP612678 | snp | C | T | V11V | IMV heparin binding surface protein |
| 87400 | OP612674, OP612679, OP612681, OP612682, OP612683, OP612686, OP612687, OP612688 | snp; APOBEC3 | G | A | H740Y | RAP94 |
| 87467 | OP612674, OP612679, OP612681, OP612682, OP612683, OP612686, OP612687, OP612688 | snp; APOBEC3 | G | A | F717F | RAP94 |
| 88810 | OP612686 | snp; APOBEC3 | G | A | R270C | RAP94 |
| 88906 | OP612679 | snp; APOBEC3 (possible error) | C | T | E238K | RAP94 |

| | | | | | | |
|---|---|---|---|---|---|---|
| 89269 | OP612677, OP612678, OP612679, OP612684, OP612687 | del (possible error) | TA | T | F116fs | RAP94 |
| 90499 | OP612691 | snp; APOBEC3 | C | T | S22L | topoisomerase type IB |
| 91115 | OP612681 | snp; APOBEC3 | C | T | V227V | topoisomerase type IB |
| 91898 | OP612674, OP612679, OP612681, OP612682, OP612683, OP612686, OP612687, OP612688 | snp; APOBEC3 | G | A | | |
| 92223 | OP612677 | snp; APOBEC3 | C | T | S108L | bifunctional large subunit of mRNA capping enzyme protein |
| 92686 | OP612674, OP612675, OP612676, OP612677, OP612678, OP612680, OP612681, OP612682, OP612683, OP612684, OP612685, OP612686, OP612687, OP612688, OP612689, OP612690, OP612691 | snp | T | C | D262D | bifunctional large subunit of mRNA capping enzyme protein |
| 93000 | OP612684 | snp; APOBEC3 | C | T | S367F | bifunctional large subunit of mRNA capping enzyme protein |
| 93557 | OP612689, OP612690, OP612691 | snp; APOBEC3 | G | A | E553K | bifunctional large subunit of mRNA capping enzyme protein |
| 95010 | OP612674 | snp; APOBEC3 | G | A | E61K | virion core protein |
| 97011 | OP612677 | snp; APOBEC3 | G | A | D265N | NTPase |
| 98658 | OP612674, OP612675, OP612676, OP612677, OP612678, OP612680, OP612681, OP612682, OP612683, OP612684, OP612685, OP612686, OP612687, OP612688, OP612689, OP612690, OP612691 | ins | A | AG | | |
| 98896 | OP612674, OP612675, OP612676, OP612677, OP612678, OP612680, OP612681, OP612682, | snp | T | C | | |

| | | | | | | |
|---|---|---|---|---|---|---|
| | OP612683, OP612684, OP612685, OP612686, OP612687, OP612688, OP612689, OP612690, OP612691 | | | | | |
| 100731 | OP612674, OP612675, OP612677, OP612678, OP612679, OP612681, OP612682, OP612683, OP612684, OP612685, OP612686, OP612687, OP612688, OP612689, OP612690, OP612691 | snp | C | T | G59G | DNA-dependent RNA polymerase subunit rpo18 |
| 100811 | OP612679 | snp; APOBEC3 | G | A | R86K | DNA-dependent RNA polymerase subunit rpo18 |
| 101140 | OP612677 | snp | C | T | E260K | IMV membrane protein |
| 104917 | OP612680 | snp | T | C | D108G | bifunctional ATPase/nucleoside triphosphate phosphohydrolase-I |
| 105295 | OP612674, OP612675, OP612676, OP612677, OP612678, OP612680, OP612681, OP612682, OP612683, OP612684, OP612685, OP612686, OP612687, OP612688, OP612689, OP612690, OP612691 | snp | T | C | R281R | bifunctional small subunit of mRNA capping enzyme protein |
| 114010 | OP612675, OP612676, OP612678, OP612680, OP612684 | snp; APOBEC3 | C | T | G619E | 82 kDa large subunit of early gene transcription factor VETF |
| 114828 | OP612689, OP612690, OP612691 | snp | G | A | A346A | 82 kDa large subunit of early gene transcription factor VETF |
| 119465 | OP612674, OP612679, OP612681, OP612682, OP612683, OP612686, OP612687, OP612688 | snp; APOBEC3 | C | T | D98N | precursor p4a of core protein 4a |
| 119887 | OP612674, OP612675, OP612676, OP612677, OP612678, OP612680, OP612681, OP612682, OP612683, OP612684, | snp | C | T | T39T | hypothetical protein |

| | | | | | | |
|---|---|---|---|---|---|---|
| | OP612685, OP612686, OP612687, OP612688, OP612689, OP612690, OP612691 | | | | | |
| 121284 | OP612680, OP612681, OP612684, OP612689 | del (possible error) | AT | A | N6fs | core protein |
| 121489 | OP612674, OP612679, OP612681, OP612682, OP612683, OP612686, OP612687, OP612688 | snp | C | T | A17T | IMV membrane protein |
| 121623 | OP612675, OP612676, OP612678 | snp; APOBEC3 | G | A | | |
| 123169 | OP612677 | snp; APOBEC3 | C | T | E106K | soluble myristylprotein |
| 124200 | OP612689, OP612690, OP612691 | snp; APOBEC3 | G | A | D29N | DNA helicase |
| 124299 | OP612686, OP612687 | snp; APOBEC3 | G | A | E62K | DNA helicase |
| 125418 | OP612674, OP612679, OP612681, OP612682, OP612683, OP612686, OP612687, OP612688 | snp; APOBEC3 (possible error) | G | A | E435K | DNA helicase |
| 125661 | OP612677 | snp; APOBEC3 | C | T | D50N | hypothetical protein |
| 128245 | OP612674, OP612675, OP612676, OP612677, OP612678, OP612680, OP612681, OP612682, OP612683, OP612684, OP612685, OP612686, OP612687, OP612688, OP612689, OP612690, OP612691 | snp; APOBEC3 | G | A | D100N | 45 kDa large subunit of intermediate gene transcription factor VITF-3 |
| 128343 | OP612677 | snp | C | T | V132V | 45 kDa large subunit of intermediate gene transcription factor VITF-3 |
| 128867 | OP612686, OP612687 | snp; APOBEC3 | C | T | S307L | 45 kDa large subunit of intermediate gene transcription factor VITF-3 |
| 129931 | OP612674, OP612675, OP612676, OP612677, OP612678, OP612680, OP612681, OP612682, OP612683, OP612684, OP612685, OP612686, OP612687, OP612688, | snp; APOBEC3 | G | A | G280D | DNA-dependent RNA polymerase subunit rpo132 |

| | | | | | | |
|---|---|---|---|---|---|---|
| | OP612689, OP612690, OP612691 | | | | | |
| 132689 | OP612691 | snp; APOBEC3 | C | T | | |
| 133263 | OP612680 | del (possible error) | TTTTTTTTTC | T | | |
| 133265 | OP612675, OP612676 | del (possible error) | TTTTTTTC | T | | |
| 133267 | OP612686, OP612687, OP612689 | del (possible error) | TTTTTC | T | | |
| 133268 | OP612677, OP612682 | del (possible error) | TTTTC | T | | |
| 133269 | OP612678, OP612681, OP612684, OP612691 | del (possible error) | TTTC | T | | |
| 133270 | OP612685, OP612688 | del (possible error) | TTC | T | | |
| 133271 | OP612690 | del (possible error) | TC | T | | |
| 133272 | OP612679 | del (possible error) | C | T | | |
| 133272 | OP612683 | del (possible error) | C | TTT | | |
| 133272 | OP612674 | del (possible error) | C | TTTT | | |
| 133318 | OP612674, OP612675, OP612676, OP612677, OP612678, OP612680, OP612681, OP612682, OP612683, OP612684, OP612685, OP612686, OP612687, OP612688, OP612689, OP612690, OP612691 | snp | C | T | | |
| 133336 | OP612686, OP612687 | del | GCAATCTTTC T | G | | |
| 134251 | OP612677 | snp; APOBEC3 | C | T | R672K | cowpox A-type inclusion protein |
| 136733 | OP612674, OP612675, OP612676, OP612678, OP612679, OP612681, OP612682, OP612683, OP612684, OP612685, OP612686, OP612689, OP612690, OP612691 | Complex (possible error) | CATNATCATC | TATGAT | D370fs | cowpox A-type inclusion protein |

| | | | | | | |
|---|---|---|---|---|---|---|
| 136736 | OP612687 | Complex (possible error) | NATC | TATG | missense_variant c.1111_1114delGATTinsCATA p.D???371HN | cowpox A-type inclusion protein |
| 137809 | OP612674, OP612675, OP612676, OP612677, OP612678, OP612680, OP612681, OP612682, OP612683, OP612684, OP612685, OP612686, OP612687, OP612688, OP612689, OP612690, OP612691 | snp | G | A | T14M | cowpox A-type inclusion protein |
| 138649 | OP612679 | del (possible error) | TA | T | F8fs | IMV surface protein |
| 138753 | OP612681 | snp | G | A | T280I | DNA-dependent RNA polymerase rpo35 |
| 138973 | OP612679 | snp; APOBEC3 | C | T | D207N | DNA-dependent RNA polymerase rpo35 |
| 140286 | OP612674, OP612675, OP612676, OP612677, OP612678, OP612680, OP612681, OP612682, OP612683, OP612684, OP612685, OP612686, OP612687, OP612688, OP612689, OP612690, OP612691 | ins | A | AATAACAATT | N123_C124(ins)NYN | hypothetical protein |
| 141473 | OP612674, OP612675, OP612676, OP612677, OP612678, OP612680, OP612681, OP612682, OP612683, OP612684, OP612685, OP612686, OP612687, OP612688, OP612689, OP612690, OP612691 | snp; APOBEC3 | G | A | E67K | bifunctional EEV membrane phosphoglycoprotein |
| 141537 | OP612674, OP612675, OP612676, OP612677, OP612678, OP612680, OP612681, OP612682, OP612683, OP612684, OP612685, OP612686, OP612687, OP612688, | snp | C | T | A88V | bifunctional EEV membrane phosphoglycoprotein |

| | | | | | | |
|---|---|---|---|---|---|---|
| | OP612689, OP612690, OP612691 | | | | | |
| 141930 | OP612683 | snp; APOBEC3 | C | T | L36L | EEV glycoprotein |
| 142623 | OP612674, OP612675, OP612676, OP612677, OP612678, OP612680, OP612681, OP612682, OP612683, OP612684, OP612685, OP612686, OP612687, OP612688, OP612689, OP612690, OP612691 | snp | C | A | T83T | hypothetical protein |
| 144018 | OP612688 | snp; APOBEC3 | C | T | I110I | hypothetical protein |
| 144832 | OP612678 | snp; APOBEC3 | G | A | S250F | CD47-like putative membrane protein |
| 145049 | OP612689 | snp; APOBEC3 | G | A | P178S | CD47-like putative membrane protein |
| 146225 | OP612677 | snp; APOBEC3 | C | T | | |
| 146780 | OP612675, OP612676, OP612678, OP612680, OP612684 | snp; APOBEC3 | C | T | E58K | bifunctional secreted glycoprotein |
| 146860 | OP612674, OP612675, OP612676, OP612677, OP612678, OP612680, OP612681, OP612682, OP612683, OP612684, OP612685, OP612686, OP612687, OP612688, OP612689, OP612690, OP612691, | snp | T | C | D31G | bifunctional secreted glycoprotein |
| 147034 | OP612674, OP612675, OP612677, OP612678, OP612680, OP612684, OP612686, OP612688, OP612689, OP612690, OP612691 | ins | A | ATATTTTATATTTTATATTT | | |
| 148566 | OP612674, OP612679, OP612681, OP612682, OP612683, OP612686, OP612687, OP612688 | snp; APOBEC3 | G | A | I328I | bifunctional hydroxysteroid dehydrogenase |
| 148682 | OP612689, OP612690, OP612691 | del | TCATATCA | T | N287fs | bifunctional hydroxysteroid dehydrogenase |

| | | | | | | |
|---|---|---|---|---|---|---|
| 149419 | OP612689, OP612690, OP612691 | | G | A | S44L | bifunctional hydroxysteroid dehydrogenase |
| 149898 | OP612674, OP612675, OP612676, OP612677, OP612678, OP612680, OP612681, OP612682, OP612683, OP612684, OP612685, OP612686, OP612687, OP612688, OP612689, OP612690, OP612691 | snp | A | G | A101A | Cu-Zn superoxide dismutase-like protein |
| 150026 | OP612689, OP612690, OP612691 | snp; APOBEC3 | C | T | Q22* | Toll/IL1-receptor-like protein |
| 150032 | OP612674, OP612675, OP612676, OP612677, OP612678, OP612680, OP612681, OP612682, OP612683, OP612684, OP612685, OP612686, OP612687, OP612688, OP612689, OP612690, OP612691 | snp | A | G | N24D | Toll/IL1-receptor-like protein |
| 150560 | OP612682 | snp | C | T | L200L | Toll/IL1-receptor-like protein |
| 150985 | OP612677 | snp; APOBEC3 | C | T | | |
| 151204 | OP612677 | snp; APOBEC3 | C | T | | |
| 151459 | OP612677 | snp; APOBEC3 | C | T | | |
| 151775 | OP612675, OP612676 | snp | T | G | F38C | thymidylate kinase |
| 152083 | OP612683, OP612685 | snp; APOBEC3 | G | A | E141K | thymidylate kinase |
| 152585 | OP612688 | snp; APOBEC3 | G | A | | |
| 154169 | OP612689, OP612690, OP612691 | snp; APOBEC3 | G | A | D442N | DNA ligase |
| 156917 | OP612676 | del (possible error) | CA | C | | |
| 158319 | OP612674, OP612675, OP612676, OP612677, OP612678, OP612680, OP612681, OP612682, OP612683, OP612684, OP612685, OP612686, OP612687, OP612688, | ins | T | TA | | |

| | | | | | | |
|---|---|---|---|---|---|---|
| | OP612689, OP612690, OP612691 | | | | | |
| 160327 | OP612674, OP612675, OP612676, OP612677, OP612678, OP612680, OP612681, OP612682, OP612683, OP612684, OP612685, OP612686, OP612687, OP612688, OP612689, OP612690, OP612691 | snp | T | G | | |
| 160981 | OP612676 | Ins (possible error) | G | GA | S72fs | ser/thr kinase |
| 161005 | OP612674, OP612675, OP612676, OP612677, OP612678, OP612680, OP612681, OP612682, OP612683, OP612684, OP612685, OP612686, OP612687, OP612688, OP612689, OP612690, OP612691 | snp | T | C | H77H | ser/thr kinase |
| 161020 | OP612674, OP612675, OP612676, OP612677, OP612678, OP612680, OP612681, OP612682, OP612683, OP612684, OP612685, OP612686, OP612687, OP612688, OP612689, OP612690, OP612691 | snp | G | A | T82T | ser/thr kinase |
| 161769 | OP612679 | snp; APOBEC3 | C | T | F9F | hypothetical protein |
| 162106 | OP612689, OP612690, OP612691 | snp; APOBEC3 | C | T | H122Y | hypothetical protein |
| 162396 | OP612680, OP612684, OP612686, OP612687 | snp; APOBEC3 | G | A | L218L | hypothetical protein |
| 163342 | OP612674, OP612675, OP612677, OP612678, OP612679, OP612681, OP612682, OP612683, OP612684, OP612685, OP612686, OP612687, OP612688, OP612689, OP612690, OP612691 | del | TTAAC | T | | |

| | | | | | | |
|---|---|---|---|---|---|---|
| 164989 | OP612674, OP612679, OP612681, OP612682, OP612683, OP612686, OP612687, OP612688 | snp; APOBEC3 | C | T | L500L | ankyrin-like protein |
| 165845 | OP612678 | snp | G | A | P189P | EEV type-I membrane glycoprotein |
| 165939 | OP612689, OP612690, OP612691 | snp; APOBEC3 | C | T | P221S | EEV type-I membrane glycoprotein |
| 166811 | OP612674, OP612675, OP612676, OP612677, OP612678, OP612680, OP612681, OP612682, OP612683, OP612684, OP612685, OP612686, OP612687, OP612688, OP612689, OP612690, OP612691 | ins | A | ATT | Y166fs | ankyrin-like protein |
| 166941 | OP612688 | snp; APOBEC3 | C | T | S19L | bifunctional 21 kDa precursor protein of 18 kDa membrane protein |
| 167811 | OP612674, OP612675, OP612676, OP612677, OP612678, OP612680, OP612681, OP612682, OP612683, OP612684, OP612685, OP612686, OP612687, OP612688, OP612689, OP612690, OP612691 | snp | G | T | R108I | soluble interferon-gamma receptor-like protein |
| 168275 | OP612674, OP612679, OP612681, OP612682, OP612683, OP612686, OP612687, OP612688 | snp; APOBEC3 (possible error) | C | T | L263F | soluble interferon-gamma receptor-like protein |
| 168360 | OP612689, OP612690, OP612691 | snp | G | A | | |
| 168520 | OP612687 | snp; APOBEC3 | G | A | E46K | 6 kDa intracellular viral protein |
| 169876 | OP612674, OP612675, OP612677, OP612678, OP612679, OP612681, OP612683, OP612684, OP612685, OP612686, OP612689, OP612690, OP612691 | ins | T | TCAGATA | T32_D33dup | hypothetical protein |

| | | | | | | |
|---|---|---|---|---|---|---|
| 171045 | OP612689, OP612690, OP612691 | ins | C | CT | | |
| 171824 | OP612674, OP612675, OP612676, OP612677, OP612678, OP612680, OP612681, OP612682, OP612683, OP612684, OP612685, OP612686, OP612687, OP612688, OP612689, OP612690, OP612691, | snp; APOBEC3 | G | A | E230K | bifunctional SPI-2/CrmA protein/IL-1 convertase |
| 174703 | OP612674, OP612675, OP612676, OP612677, OP612678, OP612681, OP612682, OP612683, OP612684, OP612685, OP612686, OP612687, OP612688, OP612691 | ins | T | TGATGAA | | |
| 177899 | OP612689, OP612690, OP612691 | snp; APOBEC3 | C | T | R689C | ankyrin-like protein |
| 178219 | OP612674, OP612675, OP612676, OP612677, OP612678, OP612681, OP612682, OP612683, OP612684, OP612685, OP612686, OP612687, OP612688, OP612689, OP612690, OP612691 | del | GTTT | G | | |
| 178605 | OP612674, OP612675, OP612677, OP612678, OP612679, OP612681, OP612682, OP612683, OP612684, OP612685, OP612686, OP612687, OP612688, OP612689, OP612690, OP612691 | snp; APOBEC3 | G | A | | |
| 180838 | OP612684 | snp; APOBEC3 | G | A | D47N | hypothetical protein |
| 181427 | OP612674, OP612675, OP612677, OP612678, OP612679, OP612681, OP612682, OP612683, OP612684, OP612685, OP612686, OP612687, | ins | G | GA | | |

| | | | | | | |
|---|---|---|---|---|---|---|
| | OP612688, OP612689, OP612690, OP612691 | | | | | |
| 182377 | OP612682 | snp; APOBEC3 | G | A | D281N | putative membrane-associated glycoprotein |
| 184218 | OP612688 | snp | G | T | T894T | putative membrane-associated glycoprotein |
| 184716 | OP612688 | snp; APOBEC3 | C | T | I1060I | putative membrane-associated glycoprotein |
| 186604 | OP612684 | snp | G | A | E1690K | putative membrane-associated glycoprotein |
| 187253 | OP612674, OP612675, OP612677, OP612682, OP612683, OP612685, OP612686, OP612687, OP612688, OP612689, OP612690, OP612691 | ins | A | AT | | |
| 187608 | OP612674, OP612677, OP612679, OP612681, OP612682, OP612685, OP612686, OP612687, OP612688 | snp; APOBEC3 | C | T | | |
| 188683 | OP612674 | snp; APOBEC3 | C | T | | |
| 189069 | OP612677 | snp; APOBEC3 | G | A | | |

Mutations in 18 MPXV genomes isolated from humans in Nigeria between January 2019 and January 2020, relative to the 1971 zoonotic Nigeria isolate KJ642617.

**Table S2.**

| Statistic \ Temporal group | 2018 | 2019 | 2021 | 2022 | 2022 (Other) |
|---|---|---|---|---|---|
| **n** | 2 | 18 | 2 | 288 | 2 |
| **Minimum** | 0 | 2 | 31 | 34 | 27 |
| **Lower (25$^{th}$) Quartile** | 0.25 | 4 | 31.8 | 41 | 28 |
| **Median** | 0.5 | 8.5 | 32.5 | 42 | 29 |
| **Upper (75$^{th}$) Quartile** | 0.75 | 12.8 | 33.2 | 43 | 30 |
| **Maximum** | 1 | 20 | 34 | 48 | 31 |
| **Kruskall-Wallis test for 2019 vs 2022** | | | | $P$=8.37x10$^{-14}$ | |

Summary statistics for APOBEC3 style mutation frequency by temporal group against a baseline of 2018 UK isolates, and statistical analysis of 2019 values vs 2022.

**Table S3.**

| Statistic \ Temporal group | 2017 | 2018 | 2019 | 2021 | 2022 | 2022 (Other) |
|---|---|---|---|---|---|---|
| **n** | 7 | 10 | 18 | 2 | 288 | 2 |
| **Minimum** | 15 | 13 | 18 | 41 | 58 | 37 |
| **Lower (25$^{th}$) Quartile** | 15.5 | 14.2 | 23 | 45.8 | 66 | 38.2 |
| **Median** | 16 | 22.5 | 25 | 50.5 | 67 | 39.5 |
| **Upper (75$^{th}$) Quartile** | 20.5 | 25 | 27.5 | 55.2 | 68 | 40.8 |
| **Maximum** | 28 | 26 | 33 | 60 | 72 | 42 |

Summary statistics for APOBEC3 style mutation frequency by temporal group against a baseline of 1971 Zoonotic Nigeria isolate KJ642617.

**Table S4.**

| | 2017 | 2018 | 2019 |
|---|---|---|---|
| **2017** | | | |
| **2018** | >0.999 | | |
| **2019** | >0.999 | >0.999 | |
| **2022** | 9.26e-06 | 1.15e-07 | 1.22e-11 |

*P* values from Dunn's test for pairwise comparisons of APOBEC3 style mutation frequency by temporal group against a baseline of 1971 Zoonotic Nigeria isolate KJ642617. Bonferroni correction was applied to adjust for multiple testing, following Kruskal-Wallis test result indicating significant differences between groups ($p<0.001$).

**Table S5.**

| Genome identifier | APOBEC3 count vs MT903345 | APOBEC3 count vs KJ642617 | Year of isolation | Country |
|---|---|---|---|---|
| KJ642617 | ND | N/A | 1971 | Nigeria |
| MK783027.1 | ND | 15 | 2017 | Nigeria |
| MK783028.1 | ND | 15 | 2017 | Nigeria |
| MK783029.1 | ND | 16 | 2017 | Nigeria |
| MK783030.1 | ND | 21 | 2017 | Nigeria |
| MK783031.1 | ND | 16 | 2017 | Nigeria |
| MK783032.1 | ND | 20 | 2017 | Nigeria |
| MK783033.1 | ND | 28 | 2017 | Nigeria |
| MN648051.1 | ND | 25 | 2018 | Israel |
| OP612674 | 6 | 29 | 2019 | Nigeria |
| OP612675 | 12 | 22 | 2019 | Nigeria |
| OP612676 | 10 | 20 | 2019 | Nigeria |
| OP612677 | 20 | 31 | 2019 | Nigeria |
| OP612678 | 13 | 23 | 2019 | Nigeria |
| OP612679 | 8 | 33 | 2019 | Nigeria |
| OP612680 | 12 | 23 | 2019 | Nigeria |
| OP612681 | 7 | 29 | 2019 | Nigeria |
| OP612682 | 6 | 28 | 2019 | Nigeria |
| OP612683 | 2 | 26 | 2019 | Nigeria |
| OP612684 | 12 | 23 | 2019 | Nigeria |
| OP612685 | 2 | 24 | 2019 | Nigeria |
| OP612686 | 8 | 33 | 2019 | Nigeria |
| OP612687 | 10 | 35 | 2019 | Nigeria |
| OP612688 | 11 | 33 | 2019 | Nigeria |
| OP612689 | 14 | 24 | 2019 | Nigeria |
| OP612690 | 13 | 24 | 2019 | Nigeria |
| OP612691 | 16 | 26 | 2020 | Nigeria |
| MT903337.1 | ND | 13 | 2018 | Nigeria |
| MT903338.1 | ND | 14 | 2018 | Nigeria |
| MT903339.1 | ND | 14 | 2018 | Nigeria |
| MT903340.1 | ND | 15 | 2018 | Nigeria |
| MT903341.1 | ND | 22 | 2018 | Nigeria |
| MT903342.1 | ND | 26 | 2018 | Nigeria |
| MT903343.1 | ND | 23 | 2018 | UK |
| MT903344.1 | ND | 25 | 2018 | UK |
| MT903345.1 | N/A | 25 | 2018 | UK |
| ON563414.3 | 41 | 66 | 2022 | USA |

| | | | | |
|---|---|---|---|---|
| ON568298.1 | 42 | 68 | 2022 | Germany |
| ON585029.1 | 37 | 59 | 2022 | Portugal |
| ON585030.1 | 41 | 65 | 2022 | Portugal |
| ON585031.1 | 41 | 65 | 2022 | Portugal |
| ON585032.1 | 41 | 66 | 2022 | Portugal |
| ON585033.1 | 42 | 67 | 2022 | Portugal |
| ON585034.1 | 41 | 66 | 2022 | Portugal |
| ON585035.1 | 41 | 66 | 2022 | Portugal |
| ON585036.1 | 41 | 64 | 2022 | Portugal |
| ON585037.1 | 44 | 69 | 2022 | Portugal |
| ON585038.1 | 44 | 69 | 2022 | Portugal |
| ON595760.2 | 41 | 66 | 2022 | Switzerland |
| ON602722.2 | 40 | 65 | 2022 | France |
| ON609725.2 | 45 | 70 | 2022 | Spain |
| ON614676.1 | 36 | 61 | 2022 | Italy |
| ON615424.1 | 43 | 69 | 2022 | Netherlands |
| ON619835.2 | 41 | 66 | 2022 | UK |
| ON619836.2 | 43 | 68 | 2022 | UK |
| ON619837.2 | 42 | 67 | 2022 | UK |
| ON619838.2 | 43 | 68 | 2022 | UK |
| ON622712.1 | 42 | 67 | 2022 | Belgium |
| ON622713.1 | 43 | 68 | 2022 | Belgium |
| ON622718.1 | 44 | 69 | 2022 | Spain |
| ON622720.1 | 39 | 60 | 2022 | Switzerland |
| ON622721.1 | 44 | 67 | 2022 | Italy |
| ON622722.2 | 44 | 69 | 2022 | France |
| ON627808.1 | 41 | 67 | 2022 | USA |
| ON631241.1 | 41 | 66 | 2022 | Slovenia |
| ON631963.1 | 43 | 69 | 2022 | Australia |
| ON637938.1 | 42 | 67 | 2022 | Germany |
| ON637939.1 | 43 | 68 | 2022 | Germany |
| ON644344.1 | 42 | 67 | 2022 | Italy |
| ON649708.1 | 41 | 66 | 2022 | Portugal |
| ON649709.1 | 41 | 66 | 2022 | Portugal |
| ON649710.1 | 41 | 65 | 2022 | Portugal |
| ON649711.1 | 41 | 65 | 2022 | Portugal |
| ON649712.1 | 41 | 66 | 2022 | Portugal |
| ON649713.1 | 44 | 69 | 2022 | Portugal |
| ON649714.1 | 41 | 65 | 2022 | Portugal |
| ON649715.1 | 41 | 65 | 2022 | Portugal |
| ON649716.1 | 42 | 66 | 2022 | Portugal |

| | | | | |
|---|---|---|---|---|
| ON649717.1 | 42 | 67 | 2022 | Portugal |
| ON649718.1 | 41 | 66 | 2022 | Portugal |
| ON649719.1 | 41 | 66 | 2022 | Portugal |
| ON649720.1 | 41 | 66 | 2022 | Portugal |
| ON649721.1 | 41 | 66 | 2022 | Portugal |
| ON649722.1 | 41 | 66 | 2022 | Portugal |
| ON649723.1 | 41 | 66 | 2022 | Portugal |
| ON649724.1 | 41 | 66 | 2022 | Portugal |
| ON649725.1 | 41 | 66 | 2022 | Portugal |
| ON649879.1 | 41 | 66 | 2022 | Israel |
| ON674051.1 | 27 | 37 | 2022 | USA |
| ON675438.1 | 31 | 42 | 2022 | USA |
| ON676703.1 | 41 | 66 | 2022 | USA |
| ON676704.1 | 43 | 68 | 2022 | USA |
| ON676705.1 | 42 | 67 | 2022 | USA |
| ON676706.1 | 41 | 66 | 2022 | USA |
| ON676707.1 | 31 | 41 | 2022 | USA |
| ON676708.1 | 34 | 60 | 2022 | USA |
| ON682263.4 | 41 | 65 | 2022 | Germany |
| ON682264.4 | 41 | 65 | 2022 | Germany |
| ON682265.4 | 41 | 66 | 2022 | Germany |
| ON682266.2 | 43 | 68 | 2022 | Germany |
| ON682267.2 | 41 | 66 | 2022 | Germany |
| ON682268.3 | 43 | 68 | 2022 | Germany |
| ON682269.3 | 41 | 65 | 2022 | Germany |
| ON682270.2 | 41 | 66 | 2022 | Germany |
| ON694329.1 | 41 | 66 | 2022 | Germany |
| ON694330.1 | 45 | 70 | 2022 | Germany |
| ON694331.1 | 42 | 67 | 2022 | Germany |
| ON694332.1 | 41 | 66 | 2022 | Germany |
| ON694333.1 | 41 | 66 | 2022 | Germany |
| ON694334.1 | 42 | 67 | 2022 | Germany |
| ON694335.1 | 43 | 68 | 2022 | Germany |
| ON694336.1 | 41 | 66 | 2022 | Germany |
| ON694337.1 | 42 | 67 | 2022 | Germany |
| ON694338.1 | 41 | 66 | 2022 | Germany |
| ON694339.1 | 42 | 67 | 2022 | Germany |
| ON694340.1 | 41 | 66 | 2022 | Germany |
| ON694341.2 | 42 | 67 | 2022 | Germany |
| ON694342.1 | 43 | 68 | 2022 | Germany |
| ON720848.1 | 43 | 66 | 2022 | Spain |

| | | | | |
|---|---|---|---|---|
| ON720849.1 | 45 | 69 | 2022 | Spain |
| ON736420.2 | 41 | 66 | 2022 | Canada |
| ON745215.1 | 41 | 66 | 2022 | Italy |
| ON745225.1 | 42 | 67 | 2022 | Spain |
| ON751962.1 | 44 | 69 | 2022 | Portugal |
| ON754984.1 | 44 | 72 | 2022 | Slovenia |
| ON754985.1 | 44 | 69 | 2022 | Slovenia |
| ON754986.1 | 44 | 69 | 2022 | Slovenia |
| ON754987.1 | 45 | 70 | 2022 | Slovenia |
| ON754989.2 | 46 | 71 | 2022 | Canada |
| ON755039.1 | 42 | 67 | 2022 | France |
| ON755040.1 | 41 | 66 | 2022 | France |
| ON755231.1 | 43 | 68 | 2022 | Germany |
| ON755232.1 | 42 | 67 | 2022 | Germany |
| ON755233.1 | 42 | 67 | 2022 | Germany |
| ON755234.1 | 42 | 67 | 2022 | Germany |
| ON755235.1 | 42 | 67 | 2022 | Germany |
| ON755236.1 | 41 | 66 | 2022 | Germany |
| ON755237.1 | 41 | 66 | 2022 | Germany |
| ON755238.2 | 41 | 66 | 2022 | Germany |
| ON755239.2 | 44 | 69 | 2022 | Germany |
| ON755240.1 | 43 | 68 | 2022 | Germany |
| ON755241.1 | 42 | 67 | 2022 | Germany |
| ON755242.1 | 41 | 66 | 2022 | Germany |
| ON755243.1 | 43 | 68 | 2022 | Germany |
| ON755244.1 | 42 | 67 | 2022 | Germany |
| ON755245.1 | 41 | 66 | 2022 | Germany |
| ON755246.1 | 41 | 66 | 2022 | Germany |
| ON755247.1 | 43 | 68 | 2022 | Germany |
| ON755248.1 | 44 | 69 | 2022 | Germany |
| ON755249.2 | 43 | 68 | 2022 | Germany |
| ON755250.1 | 41 | 66 | 2022 | Germany |
| ON755251.2 | 45 | 70 | 2022 | Germany |
| ON755252.1 | 42 | 67 | 2022 | Germany |
| ON755253.1 | 41 | 66 | 2022 | Germany |
| ON755254.1 | 43 | 68 | 2022 | Germany |
| ON755255.2 | 43 | 68 | 2022 | Germany |
| ON755256.1 | 44 | 69 | 2022 | Germany |
| ON780016.1 | 41 | 67 | 2022 | Italy |
| ON780017.1 | 42 | 68 | 2022 | Italy |
| ON782021.1 | 44 | 69 | 2022 | Finland |

| | | | | |
|---|---|---|---|---|
| ON782022.1 | 41 | 66 | 2022 | Finland |
| ON782054.1 | 34 | 58 | 2022 | Spain |
| ON782055.1 | 44 | 69 | 2022 | Spain |
| ON792320.1 | 45 | 69 | 2022 | Switzerland |
| ON792321.1 | 41 | 65 | 2022 | Switzerland |
| ON792322.1 | 42 | 67 | 2022 | Switzerland |
| ON803413.1 | 46 | 71 | 2022 | Canada |
| ON803414.1 | 46 | 71 | 2022 | Canada |
| ON803415.1 | 46 | 70 | 2022 | Canada |
| ON803416.1 | 41 | 65 | 2022 | Canada |
| ON803417.1 | 42 | 67 | 2022 | Canada |
| ON803418.1 | 42 | 67 | 2022 | Canada |
| ON803419.1 | 41 | 66 | 2022 | Canada |
| ON803420.1 | 46 | 71 | 2022 | Canada |
| ON803421.1 | 41 | 66 | 2022 | Canada |
| ON803422.1 | 43 | 68 | 2022 | Canada |
| ON803423.1 | 43 | 68 | 2022 | Canada |
| ON803424.1 | 41 | 66 | 2022 | Canada |
| ON803425.1 | 41 | 66 | 2022 | Canada |
| ON803426.1 | 41 | 66 | 2022 | Canada |
| ON803427.1 | 41 | 66 | 2022 | Canada |
| ON803428.1 | 41 | 65 | 2022 | Canada |
| ON803429.1 | 41 | 66 | 2022 | Canada |
| ON803430.1 | 41 | 66 | 2022 | Canada |
| ON803431.1 | 42 | 67 | 2022 | Canada |
| ON803432.1 | 42 | 67 | 2022 | Canada |
| ON803433.1 | 41 | 65 | 2022 | Canada |
| ON803434.1 | 41 | 65 | 2022 | Canada |
| ON803435.1 | 43 | 68 | 2022 | Canada |
| ON803436.1 | 41 | 66 | 2022 | Canada |
| ON803437.1 | 41 | 66 | 2022 | Canada |
| ON803438.1 | 41 | 66 | 2022 | Canada |
| ON803439.1 | 46 | 71 | 2022 | Canada |
| ON803440.1 | 41 | 66 | 2022 | Canada |
| ON803441.1 | 41 | 66 | 2022 | Canada |
| ON803442.1 | 41 | 66 | 2022 | Canada |
| ON803443.1 | 41 | 65 | 2022 | Canada |
| ON803444.1 | 42 | 67 | 2022 | Canada |
| ON808413.1 | 42 | 67 | 2022 | UK |
| ON808414.1 | 41 | 66 | 2022 | UK |
| ON808415.1 | 41 | 66 | 2022 | UK |

| | | | | |
|---|---|---|---|---|
| ON808416.1 | 42 | 67 | 2022 | UK |
| ON808417.1 | 41 | 66 | 2022 | UK |
| ON813251.2 | 42 | 66 | 2022 | Germany |
| ON813252.2 | 41 | 66 | 2022 | Germany |
| ON813253.2 | 42 | 67 | 2022 | Germany |
| ON813254.2 | 43 | 68 | 2022 | Germany |
| ON813255.2 | 41 | 66 | 2022 | Germany |
| ON813256.2 | 42 | 67 | 2022 | Germany |
| ON813257.2 | 44 | 69 | 2022 | Germany |
| ON813258.2 | 42 | 67 | 2022 | Germany |
| ON813259.2 | 43 | 68 | 2022 | Germany |
| ON813260.2 | 44 | 69 | 2022 | Germany |
| ON813261.2 | 44 | 69 | 2022 | Germany |
| ON813262.2 | 42 | 67 | 2022 | Germany |
| ON813263.2 | 43 | 68 | 2022 | Germany |
| ON813264.2 | 42 | 67 | 2022 | Germany |
| ON813265.2 | 41 | 66 | 2022 | Germany |
| ON813266.2 | 42 | 67 | 2022 | Germany |
| ON813267.2 | 41 | 66 | 2022 | Germany |
| ON838178.1 | 44 | 69 | 2022 | Portugal |
| ON838939.1 | 43 | 68 | 2022 | Spain |
| ON838940.1 | 41 | 66 | 2022 | Spain |
| ON843163.1 | 41 | 64 | 2022 | Portugal |
| ON843164.1 | 41 | 67 | 2022 | Portugal |
| ON843165.1 | 42 | 67 | 2022 | Portugal |
| ON843166.1 | 43 | 68 | 2022 | Portugal |
| ON843167.1 | 41 | 66 | 2022 | Portugal |
| ON843168.1 | 44 | 69 | 2022 | Portugal |
| ON843169.1 | 41 | 66 | 2022 | Portugal |
| ON843170.1 | 42 | 67 | 2022 | Portugal |
| ON843171.1 | 41 | 64 | 2022 | Portugal |
| ON843172.1 | 41 | 66 | 2022 | Portugal |
| ON843173.1 | 42 | 67 | 2022 | Portugal |
| ON843174.1 | 42 | 67 | 2022 | Portugal |
| ON843175.1 | 42 | 66 | 2022 | Portugal |
| ON843176.1 | 41 | 66 | 2022 | Portugal |
| ON843177.1 | 41 | 66 | 2022 | Portugal |
| ON843178.1 | 41 | 66 | 2022 | Portugal |
| ON843179.1 | 42 | 65 | 2022 | Portugal |
| ON843180.1 | 41 | 64 | 2022 | Portugal |
| ON843181.1 | 45 | 69 | 2022 | Portugal |

| | | | | |
|---|---|---|---|---|
| ON843182.1 | 42 | 66 | 2022 | Portugal |
| ON853649.1 | 45 | 70 | 2022 | Germany |
| ON853650.1 | 43 | 68 | 2022 | Germany |
| ON853651.1 | 43 | 68 | 2022 | Germany |
| ON853652.1 | 42 | 67 | 2022 | Germany |
| ON853653.1 | 41 | 66 | 2022 | Germany |
| ON853654.1 | 41 | 66 | 2022 | Germany |
| ON853655.1 | 46 | 71 | 2022 | Germany |
| ON853656.1 | 44 | 69 | 2022 | Germany |
| ON853657.1 | 41 | 66 | 2022 | Germany |
| ON853658.1 | 42 | 67 | 2022 | Germany |
| ON853659.1 | 44 | 69 | 2022 | Germany |
| ON853660.1 | 44 | 69 | 2022 | Germany |
| ON853661.1 | 45 | 70 | 2022 | Germany |
| ON853662.1 | 44 | 69 | 2022 | Germany |
| ON853663.1 | 42 | 67 | 2022 | Germany |
| ON853664.1 | 43 | 68 | 2022 | Germany |
| ON853665.1 | 44 | 69 | 2022 | Germany |
| ON853666.1 | 42 | 67 | 2022 | Germany |
| ON853667.1 | 43 | 68 | 2022 | Germany |
| ON853668.1 | 45 | 70 | 2022 | Germany |
| ON853669.1 | 44 | 69 | 2022 | Germany |
| ON853670.1 | 44 | 69 | 2022 | Germany |
| ON853671.1 | 42 | 67 | 2022 | Germany |
| ON853672.1 | 44 | 69 | 2022 | Germany |
| ON853673.1 | 41 | 66 | 2022 | Germany |
| ON853674.1 | 41 | 66 | 2022 | Germany |
| ON853675.1 | 41 | 66 | 2022 | Germany |
| ON853676.1 | 44 | 69 | 2022 | Germany |
| ON853677.1 | 42 | 67 | 2022 | Germany |
| ON853678.1 | 42 | 67 | 2022 | Germany |
| ON853679.1 | 41 | 66 | 2022 | Germany |
| ON853680.1 | 42 | 67 | 2022 | Germany |
| ON853681.1 | 42 | 67 | 2022 | Germany |
| ON853682.1 | 42 | 67 | 2022 | Germany |
| ON872184.1 | 42 | 67 | 2022 | Ireland |
| ON880413.1 | 45 | 70 | 2022 | Brazil |
| ON880419.1 | 41 | 66 | 2022 | Belgium |
| ON880420.1 | 41 | 66 | 2022 | Belgium |
| ON880421.1 | 41 | 66 | 2022 | Belgium |
| ON880422.1 | 41 | 66 | 2022 | Belgium |

| | | | | |
|---|---|---|---|---|
| ON880505.1 | 46 | 70 | 2022 | Canada |
| ON880506.1 | 48 | 72 | 2022 | Canada |
| ON880507.1 | 46 | 69 | 2022 | Canada |
| ON880508.1 | 41 | 66 | 2022 | Canada |
| ON880509.1 | 41 | 65 | 2022 | Canada |
| ON880510.1 | 46 | 70 | 2022 | Canada |
| ON880511.1 | 45 | 70 | 2022 | Canada |
| ON880512.1 | 41 | 66 | 2022 | Canada |
| ON880513.1 | 46 | 71 | 2022 | Canada |
| ON880514.1 | 46 | 70 | 2022 | Canada |
| ON880515.1 | 46 | 70 | 2022 | Canada |
| ON880516.1 | 41 | 66 | 2022 | Canada |
| ON880517.1 | 42 | 67 | 2022 | Canada |
| ON880518.1 | 42 | 65 | 2022 | Canada |
| ON880519.1 | 41 | 62 | 2022 | Canada |
| ON880520.1 | 42 | 66 | 2022 | Canada |
| ON880521.1 | 41 | 66 | 2022 | Canada |
| ON880522.1 | 41 | 66 | 2022 | Canada |
| ON880523.1 | 46 | 71 | 2022 | Canada |
| ON880524.1 | 38 | 61 | 2022 | Canada |
| ON880525.1 | 41 | 64 | 2022 | Canada |
| ON880526.1 | 41 | 66 | 2022 | Canada |
| ON880527.1 | 41 | 66 | 2022 | Canada |
| ON880528.1 | 44 | 68 | 2022 | Canada |
| ON880529.1 | 42 | 65 | 2022 | Canada |
| ON880530.1 | 41 | 66 | 2022 | Canada |
| ON880531.1 | 41 | 65 | 2022 | Canada |
| ON880532.1 | 41 | 65 | 2022 | Canada |
| ON880533.1 | 44 | 67 | 2022 | Canada |
| ON880534.1 | 44 | 69 | 2022 | Canada |
| ON880535.1 | 41 | 65 | 2022 | Canada |
| ON880536.1 | 42 | 67 | 2022 | Canada |
| ON880537.1 | 42 | 67 | 2022 | Canada |
| ON880538.1 | 44 | 68 | 2022 | Canada |
| ON880539.1 | 41 | 66 | 2022 | Canada |
| ON880540.1 | 44 | 69 | 2022 | Canada |
| ON880541.1 | 39 | 64 | 2022 | Canada |
| ON880542.1 | 41 | 66 | 2022 | Canada |
| ON880543.1 | 44 | 69 | 2022 | Canada |
| ON880544.1 | 43 | 67 | 2022 | Canada |
| ON880545.1 | 42 | 67 | 2022 | Canada |

| | | | | |
|---|---|---|---|---|
| ON880546.1 | 41 | 66 | 2022 | Canada |
| ON880547.1 | 45 | 70 | 2022 | Canada |
| ON880548.1 | 43 | 68 | 2022 | Canada |
| ON880549.1 | 44 | 69 | 2022 | Canada |

Genomes used in this analysis, with APOBEC3 counts relative to MT903345 (only genomes from 2019 onwards) and KJ64261 (all genomes).
